# Genomic signatures of coral adaptation and recovery following a mass mortality event

**DOI:** 10.1101/2024.09.23.614560

**Authors:** James Fifer, Kelly E. Speare, Sarah E. Leinbach, Stephanie F. Hendricks, Sarah W. Davies, Noah H. Rose, Deron E. Burkepile, Thomas C. Adam, Gretchen E. Hofmann, Marie E. Strader

## Abstract

Globally, corals face an increased frequency of mass mortality events (MMEs) as populations experience repeated marine heatwaves which disrupt their obligate algal symbiosis. Despite greater occurrences of MMEs, the relative roles of the environment, host, and symbiont genetic variation in survival and recovery remain unresolved. We explore these roles using high-resolution temporal and spatial whole genome sequencing of corals before, after, and several years following a MME. We show that host genetics has an impact on bleaching and mortality and that selected alleles important for adaptation persist through the next generation. We also demonstrate that survival for the bleaching event was highly polygenic and that allele frequency shifts have reef habitat specificity. This study reveals how MMEs reshape the genomic landscape and the spatial and temporal distribution of genomic diversity within coral populations facing severe threats from global change.

**SIGNIFICANCE:** Investigations of natural selection in coral genomes following bleaching events have primarily relied on indirect inference, using contemporary populations to explore signatures of past selective pressures, or failed to link these events to the subsequent generation. This has left an open question about the ability of a coral population to adapt from host standing genetic variation in the face of a bleaching event. We demonstrate rapid evolution in a Mo’orean coral population following an MME from a marine heatwave by capturing allele frequency shifts through time. However, the complex polygenic architecture of bleaching survival shows strong habitat specificity, complicating the path towards a genomic prediction of bleaching.

## INTRO

As a consequence of anthropogenic global change, mass mortality events (MMEs) are increasing in both frequency and magnitude across diverse marine ecosystems (Fey et al., 2015). Rapid decreases in population size are theorized to lower genetic diversity (Nei et al., 1975) and thus reduce the variation upon which selection acts, constraining the adaptability of survivors to future stressors (Du et al., 2016; Radwan et al., 2010). Despite this, MME-driven reductions in genetic diversity have limited empirical support in marine systems (but see Gurgel et al. (2020)). It is unclear if this is due to temporal limitations of many MME genetic diversity studies (*e.g.,* sampling only one generation following an MME) or rooted in population dynamics that protect against a loss of genetic diversity (*e.g.,* large population sizes; Nei et al. (1975)). MMEs can also lead to selection for advantageous alleles (*e.g.,* Auteri & Knowles, 2020; Campbell-Staton et al., 2017; Gignoux-Wolfsohn et al., 2021; Holland et al., 2022; Schiebelhut et al., 2018), suggesting population persistence by survivors may allow future generations to be better equipped against stressors similar to those that triggered the MME. As MMEs shape the evolutionary direction of populations and species, their increased prevalence presents a critical concern for the long-term viability and resilience of marine ecosystems.

Coral reefs, which are responsible for hosting over a quarter of the ocean’s total biodiversity (Knowlton et al., 2010), have faced numerous MMEs globally in recent decades due to sea surface temperatures exceeding their physiological limits (Eakin et al., 2019). MMEs in reef-building corals typically result from marine heatwaves (MHWs) that cause coral bleaching, the breakdown of the symbiosis between the coral and its endosymbiotic microalgal partner (family Symbiodiniaceae) (Douglas, 2003). Despite corals’ vulnerability to increased temperatures, variation in response to heat stress is observed within and among coral species (Dixon et al., 2015; Loya et al., 2001). While this variation can be at least partially attributed to the genera or species of symbiont hosted (*e.g.,* Baker et al., 2004; Berkelmans & Van Oppen, 2006; Jones et al., 2008; Kemp et al., 2023) and microenvironment differences (Fabricius, 2006), there is evidence for a heritable host basis for heat stress resilience (Quigley & van Oppen, 2022; and reviewed in Bairos-Novak et al., 2021; Howells et al., 2022), suggesting there is variation in heat stress tolerance for selection to act upon during MMEs.

How fast a species can adapt to novel environments from standing within-species genetic variation is an outstanding question in evolutionary ecology and critically time-sensitive, as organisms face rapidly changing environments globally. It has been theorized that long-term reef resilience relies on natural selection favoring heat-tolerant alleles that exist in the metapopulation prior to warming (Matz et al., 2020). While modern DNA sequences have been used to successfully identify loci that have evolved in response to natural selection in corals (Cooke et al., 2020; Rose et al., 2021; Smith et al., 2022; Zhang et al., 2022), these studies are inherently limited because they investigate selection indirectly, finding its signature in nearby neutral variation. Temporal sampling across an MME allows for a direct examination of loci under selection, but to date coral genomic research has focused on a single timepoint of an MME (*e.g.,* bleaching timepoint; (Fuller et al., 2020)) or used too few molecular markers and/or individuals (Thomas et al., 2024) to capture evolutionary changes.

In 2019, a prolonged MHW occurred around the island of Mo’orea, French Polynesia and the major reef-building genus of corals, *Acropora*, experienced 50-80% mortality as a consequence of severe, prolonged bleaching (Speare et al., 2022). We performed whole genome sequencing on *Acropora hyacinthus* colonies from multiple sites around Mo’orea: 1) during the peak of the bleaching event in May 2019 before mortality, 2) after bleaching and thermal stress had subsided in October 2019, and 3) in November 2021, two years after the MME, when new recruits naïve to the 2019 bleaching event were collected. High-resolution sampling of the population before and after the mortality event and in the subsequent generation allowed us to assess temporal genomic signatures of bleaching resistance (*i.e.* avoiding bleaching) and to answer the following questions: 1) Can host genetics explain survival from a bleaching event and what is the genetic architecture of the trait? 2) Are adaptive alleles maintained in the following generation? 3) Does adaptive genetic variation differ between discrete reef habitats?

## MAIN

*Acropora hyacinthus* colonies were sampled around the island of Mo’orea as follows: 1) adults during the May 2019 bleaching event (hereafter, pre-mortality), 2) adults after bleaching had subsided in October 2019 (hereafter, post-mortality), and 3) juveniles (colonies < 8 cm long diameter) in November 2021 (hereafter, juveniles). For each timepoint, colonies were sampled from four locations around the island that were directly adjacent to Mo’orea Coral Reef Long-Term Ecological Research (MCR LTER) sites: sites 1 and 2 on the north shore, site 3 on the east shore, and site 5 on the west shore (Fig. 1B). At each MCR LTER location, colonies were sampled from at least one of three reef habitats (backreef, shallow forereef, and deep forereef), corresponding to three different depths (1-3 m, 3-5 m, 10-14 m, respectively). While adults from May 2019 and October 2019 included samples at all three reef habitats, due to weather constraints juveniles from November 2021 were only sampled at the deep forereef (Supp. Table 1). Hereafter, each combination of three reef habitats and an associated LTER location is referred to as a “site.”

**Figure 1.**
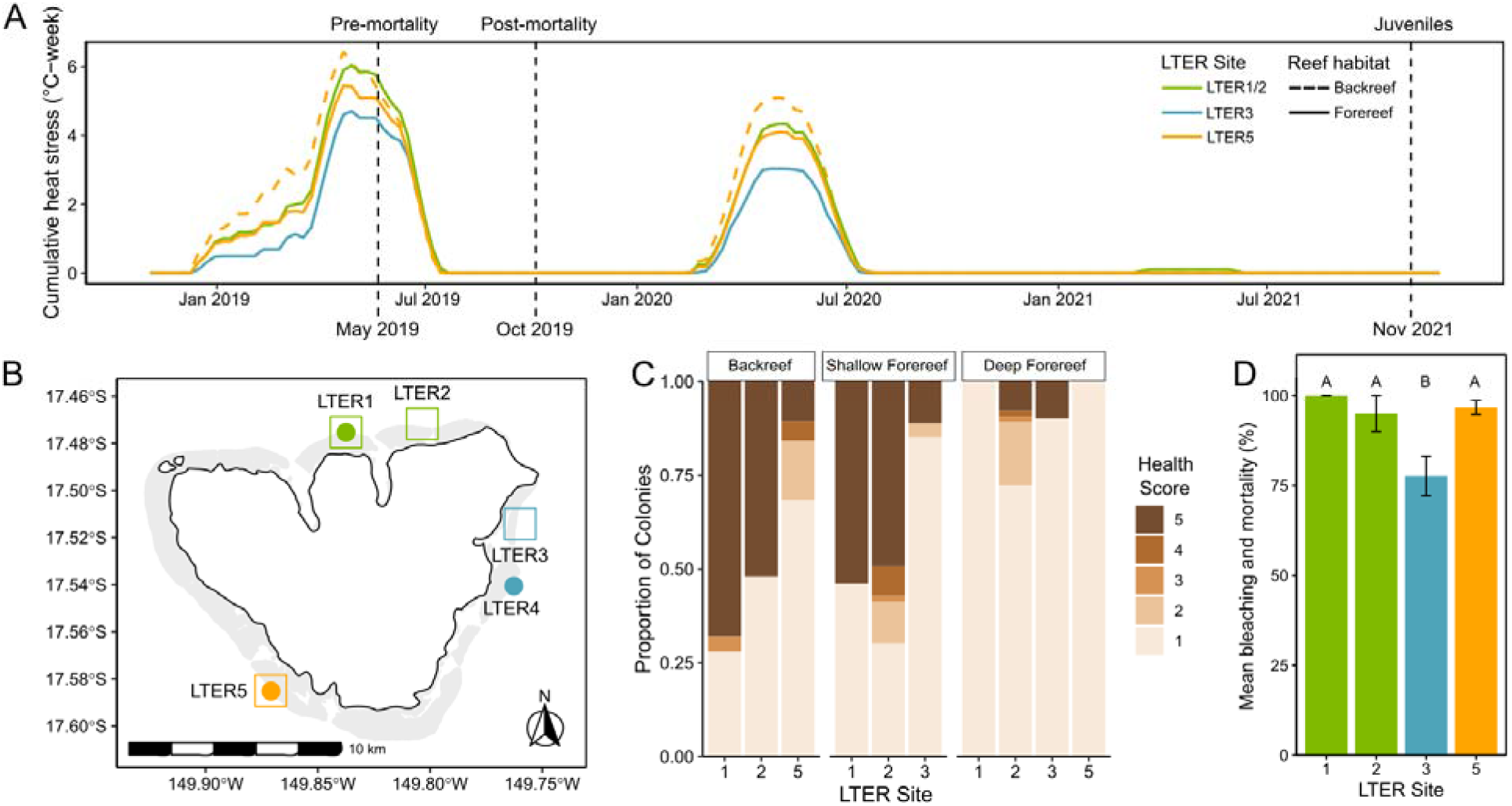
Pervasive bleaching and mortality following an extreme thermal anomaly. A) Cumulative heat stress across space and time. Vertical lines denote sampling points for sequencing. B) Mo’orea, French Polynesia. Circles represent locations where temperature data were collected; squares represent locations where samples were collected for sequencing. C) Bleaching severity observed across reef habitats and sites. A score of 1 indicates complete bleaching and 5 indicates no pigmentation loss (example photos in Supp. Fig. 12). D) Combined bleaching and mortality data from surveys in July 2019 at deep forereef locations. Letters indicate significance per pairwise Wilcoxon rank sum test.

### 2019 heatwave triggers mass bleaching and mortality

In 2019, Mo’orea experienced a prolonged MHW, with some sites experiencing 6 weeks of cumulative heat stress exceeding 29°C (Fig. 1A), a noted bleaching threshold for most corals in Mo’orea (Pratchett et al., 2013). This was the largest thermal stress event recorded in Mo’orea in the last 16 years (Supp. Fig. 1). The northern (LTERs 1 and 2) and western (LTER 5) regions experienced the largest cumulative heat stress (Fig. 1A) and subsequently displayed higher *A. hyacinthus* bleaching at the deep forereef in May 2019 (Fig. 1C) and higher bleaching and mortality in July 2019 (*p* < 0.01; Fig. 1D) compared to the eastern side of the island (LTER 3). *A. hyacinthus* experienced bleaching island-wide, with the deep forereef habitat showing the highest proportion of bleaching colonies and LTERs 1 and 5 from the deep forereef exhibiting severe paling in all colonies surveyed (Fig. 1C). As a result of this heat wave, *Acropora spp*. exhibited the highest decrease in percent cover since the 2006-2009 outbreaks of the corallivorous seastar *Acanthaster planci* (Kayal et al., 2012; Supp. Fig. 2).

### Low genetic structure in A. hyacinthus Mo’orea populations

Prior to the heat-induced mortality event in 2019, backreef and forereef sites around the island of Mo’orea showed evidence of low genetic structure (genetic connectivity (*F*_ST_)) between sites ranging from 0.013-0.02; Fig. 2A; Supp. Fig. 3). Slightly higher differentiation between sites was observed when any site was compared with a backreef site (including backreef-backreef comparisons) (*F*_ST_ between sites ranging from 0.016-0.021; Supp. Fig. 3), compared to deep forereef-deep forereef comparisons (*F*_ST_ between sites ranging from 0.012-0.016; Supp. Fig. 3). Sites were significantly differentiated pre-mortality event (*p*_permanova_ < 0.001), but the variation explained by site was low (R^2^ = 0.048) (Fig. 2B). Lack of structure was also observed in admixture analyses (Supp. Fig. 4). These results mirror *A. hyacinthus* 2b-RAD-seq data for populations in Mo’orea and Tahiti collected six years prior in 2013, where pairwise *F*_ST_ ranged from 0.011 to 0.018 (Kriefall et al., 2022). All Mo’orea samples clustered with the American Samoa *A. hyacinthus* lineage HE from (Rose et al., 2021) (Supp. Fig. 5), which was the most thermally tolerant among the four cryptic *A. hyacinthus* lineages from Rose et al. (2021).

**Figure 2.**
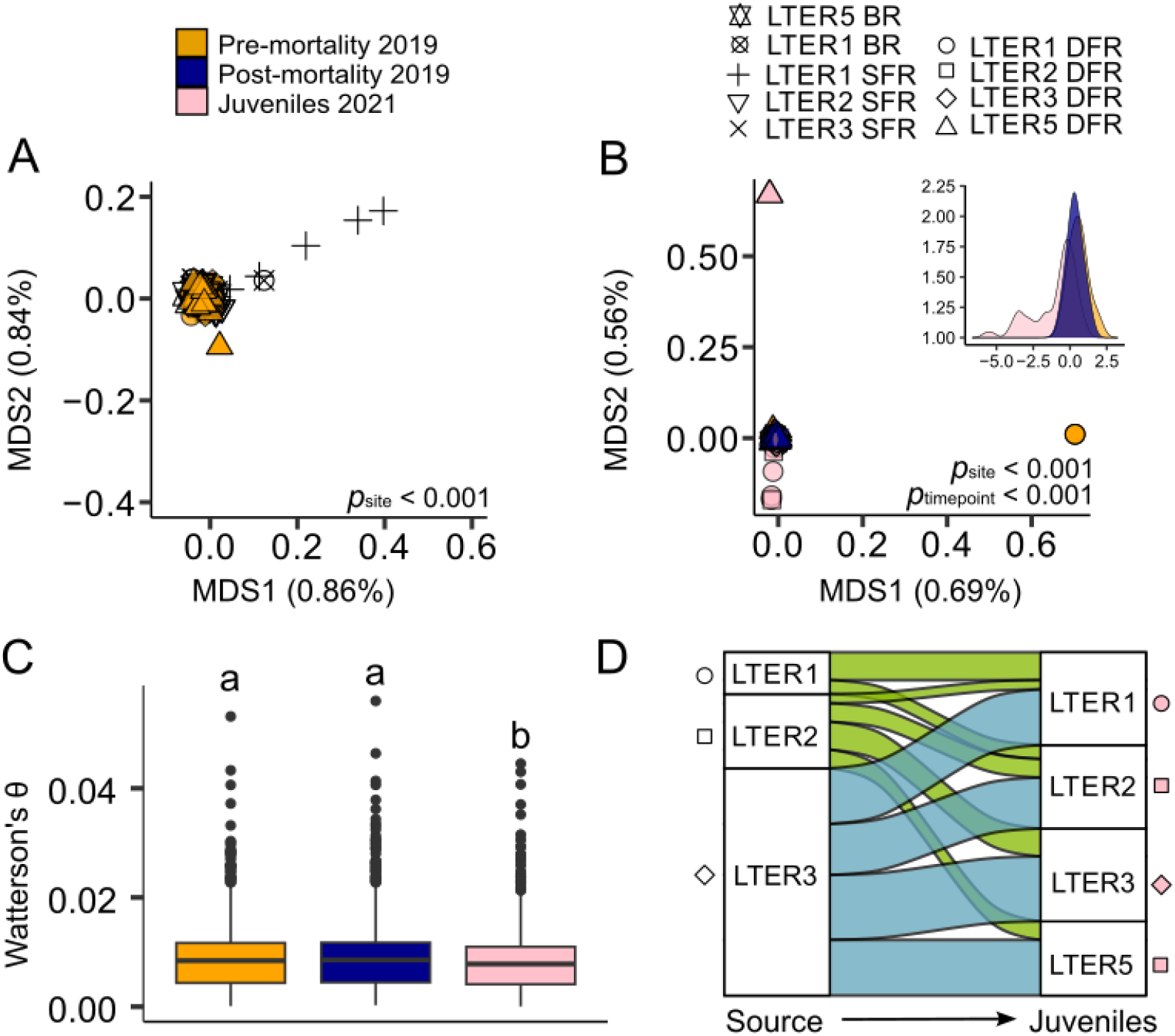
Stability of genetic diversity across adult corals and shifts in recruitment dynamics in a panmictic Mo’orean population. A) Multidimensional scaling (MDS) plot based on genetic covariance matrices of pre-mortality (May 2019; N =139) samples shows a lack of population structure. Sites that were also sampled post-mortality in 2019 and juveniles in 2021 are color filled. Sites that were exclusively sampled in May 2019 are open symbols. B) MDS plot based on genetic covariance matrices of samples from the sites that were sampled at all three timepoints and Discriminant analysis of principal components (DAPC) inset, highlighting the genome-wide similarity of samples before (N = 86), after the bleaching event (N = 71), and juveniles two years later (N = 76), except the few juveniles that cluster outside the major grouping. C) Nucleotide diversity (Watterson’s θ) between timepoints for all overlapping sites; letters denote significant differences via Dunn’s test. D) DAPC assignment between predicted source site (left column) and actual site of the juvenile (right column; N =76) based on similarity to adult samples (all 9 sites; N =244), showing the large recruitment from LTER 3 deep forereef and lack of recruitment from backreef and shallow forereef sites. BR: Backreef; SFR: Shallow forereef; DFR: Deep forereef.

### Genetic diversity and connectivity show stability through bleaching event

Genetic diversity, measured as nucleotide diversity (Watterson’s θ; Watterson, 1975), showed no difference post-mortality (October 2019) compared to pre-mortality (May 2019) (Fig. 2C), which was also true when comparing genetic diversity at the site level (Supp. Fig. 6). We similarly find no differences in pairwise θ between timepoints (Supp. Fig. 7). Individual heterozygosity did not differ post-mortality compared to pre-mortality (Supp. Fig. 8); however, at the site level, LTER 2 deep forereef exhibited decreased heterozygosity and LTER 2 shallow forereef exhibited increased heterozygosity post-mortality (Supp. Fig. 9). Empirical evidence of loss of genetic diversity directly following MMEs in marine systems is limited (Gurgel et al., 2020), with many instances of no observable decrease in genetic diversity, such as in kelp (Coleman et al., 2020; Klingbeil et al., 2022), eels (Pujolar et al., 2011) sea stars (Schiebelhut et al., 2018), mangroves (Arnaud-Haond et al., 2009), gorgonians (Pilczynska et al., 2016) and dogwhelk (Colson & Hughes, 2004). The stability of genetic diversity following MMEs has occasionally been attributed to additional recruitment between timepoints (*e.g.,* Klingbeil et al., (2022)), which is not the case in our system, as three months is not enough time for *A. hyacinthus* to recruit and develop into adults. Alternatively, populations with large effective population sizes (*N_e_*) are not expected to exhibit decreases in genetic diversity following population bottlenecks (Frankham, 1995; Franklin, 1980; Nei et al., 1975). While it is difficult to ascertain whether the population size of *A. hyacinthus* in Mo’orea is sufficiently large, we observe that historical N_e_ is an order of magnitude higher than populations of *A. hyacinthus* in Japan (Fifer et al., 2022) and several times larger than a declining *A. millepora* population in the Great Barrier Reef (GBR) (Matz et al., 2018) using the same methodology across studies (Supp. Fig. 10).

Genetic connectivity (*F*_ST_) between deep forereef sites showed stability across timepoints with no deviation in site differentiation post-mortality compared to pre-mortality or to juveniles (Supp. Fig. 11). In contrast, genetic diversity and individual heterozygosity island-wide were lower in juveniles collected two years after the MME compared to adult populations from both pre- and post-mortality timepoints (Fig. 2C; Supp. Fig. 9). This pattern is in line with DAPC juvenile assignment results (24 PCs retained per cross-validation (CV)) showcasing that putative sources for juveniles collected post-mortality event in 2021 were not evenly split between sites (χ^2^ = 48.185, *p* < 0.0001), with LTER 3 deep forereef serving as the dominant source. We also show juveniles found on the deep forereef had predicted sources from only LTERs 1, 2, and 3 deep forereef, and thus none were likely sourced from either backreef, shallow forereef, or LTER 5 deep forereef locations (Fig. 2D). Visualizations of population clustering using principal correspondence analyses (PCoA) and DAPC show that several juveniles fall outside the central genetic cluster (Fig. 2B), likely representing immigrants from other islands/reefs not sampled here.

It is difficult to determine whether the decrease of genetic diversity and individual heterozygosity in 2021 juveniles is reflective of normal recruitment patterns or a consequence of the heat stress events in 2019 and 2020, without a non-disturbance recruitment baseline to compare with. However, *A. hyacinthus* has shown a propensity to exhibit “sweepstakes reproductive success” (SRS; (Hedgecock, 1994)) dynamics, where a single recruitment year of surviving offspring comes from a subset of adults (Barfield et al., 2022). It is notable that the sites with the greatest mortality (LTERs 1 and 5) were the only sites to show a decrease in juvenile genetic diversity (Supp. Fig. 6). In contrast, the site that experienced the lowest heat stress (LTER 3; Fig. 1A), bleaching, and mortality (Fig. 1C; Fig. 1D), exhibited no decrease in juvenile diversity compared to adults and this site was also the major source of recruits in 2021. As *A. hyacinthus* colonies that escaped bleaching demonstrate higher fecundity than those that bleached and recovered in Mo’orea (Leinbach et al., 2021), it seems likely that the observed recruitment dynamics were a result of the less intense stress experienced at LTER 3, leading to both higher colony survival and fecundity compared to the other sites. While information on the hydrodynamic processes around the island is limited, it suggests the absence of oceanographic barriers from LTER 5 to the other sites, contrasting its absence as a source site for juveniles here (Fig. 2D). Leichter et al. (2013) suggests biological particles might stay in relatively close proximity to the island over periods of days to weeks, via a current that moves over the entire island in a counterclockwise fashion.

### Host shows highly polygenic response to the bleaching event

To identify major effect loci that might be driving heat stress tolerance, two different genome-wide association studies (GWAS) were performed: 1) for pre-mortality samples taken during the bleaching event (N = 172) using health score (1-5) as the trait for bleaching resistance (Fig. 1C; Supp. Fig. 12) and 2) for pre- and post-mortality samples using timepoint (May 2019 or October 2019) as a binary trait for survival (N = 231). We also repeated GWAS analyses looking at each reef-habitat subset. We determined a genome-wide significance threshold of *p* = 2.02*e*-9 for health score in the pre-mortality (May 2019) samples through shuffled permutation tests and a genome-wide significance threshold of *p* = 5.24*e*-7 for survival in the pre-mortality and post-mortality samples (Supp. Fig. 13). Using this approach, we failed to identify a single genetic locus that is significant genome-wide, with the exception of the backreef habitat subset. While the majority of our GWASs failed to detect large effect loci, this result could be expected considering thermal tolerance and bleaching has been shown to be highly polygenic (Rose et al., 2018; Fuller et al., 2020). However, it is possible that with sampling more colonies or, for the survival GWAS, monitoring colonies through the bleaching event and scoring colonies as survivors/non-survivors, we would be able to better detect large effect loci. The survival GWAS at the backreef habitat identified two potential “major effect loci” that pass our strict *p*-value threshold described above (Supp. Fig. 14). One of these loci falls near G-protein coupled receptor (GPCR) 45 (GPR45) and the other near XK Related 6 (XKR6). This particular GPCR has no described function in coral beyond its theorized role in the coral nervous system based off gene function in the human central nervous system (Marchese et al., 1999). However, GPCRs broadly are a class of important signaling receptors and are differentially expressed in a diversity of coral gene expression studies (Mason et al., 2012, 2023; Rose et al., 2015; Seneca & Palumbi, 2015) as well as theorized to modulate metabolite transfer between Symbiodiniaceae algae and the coral host cell (Traylor-Knowles et al., 2017). XKR6 is implicated in apoptotic cell ingestion (Giallourakis et al., 2006) and has no known function in coral.

Heat tolerance could alternatively be driven by many loci of small effect. There was a significant convergent shift in genome-wide allele frequencies following the mortality event between all sites except between the shallow forereef and backreef and all comparisons involving LTER 2 deep forereef (Fig. 3A). This finding demonstrates that bleaching survival is a highly polygenic trait, aligning with previous research on bleaching susceptibility in *A. hyacinthus* from American Samoa (Rose et al., 2018) and *A. millepora* in the GBR (Fuller et al., 2020). This signal of parallel polygenic selection suggests that there was largely a consistent selection pressure across sites and suggests many populations in Mo’orea share adaptive variants that are present at high frequencies (Barghi et al., 2020) or that there is background selection on shared deleterious variation (Buffalo & Coop, 2020). The lack of convergence with comparisons involving LTER 2 deep forereef can be explained by the sampling design. A complete lack of live colonies at LTER 2 in October 2019 at 10 m necessitated sampling 5 m deeper than had been sampled in May 2019 (Supp. Table 1) and thus these colonies were either representative of a different population and/or experienced differential selection pressures from the other reef sites.

**Figure 3.**
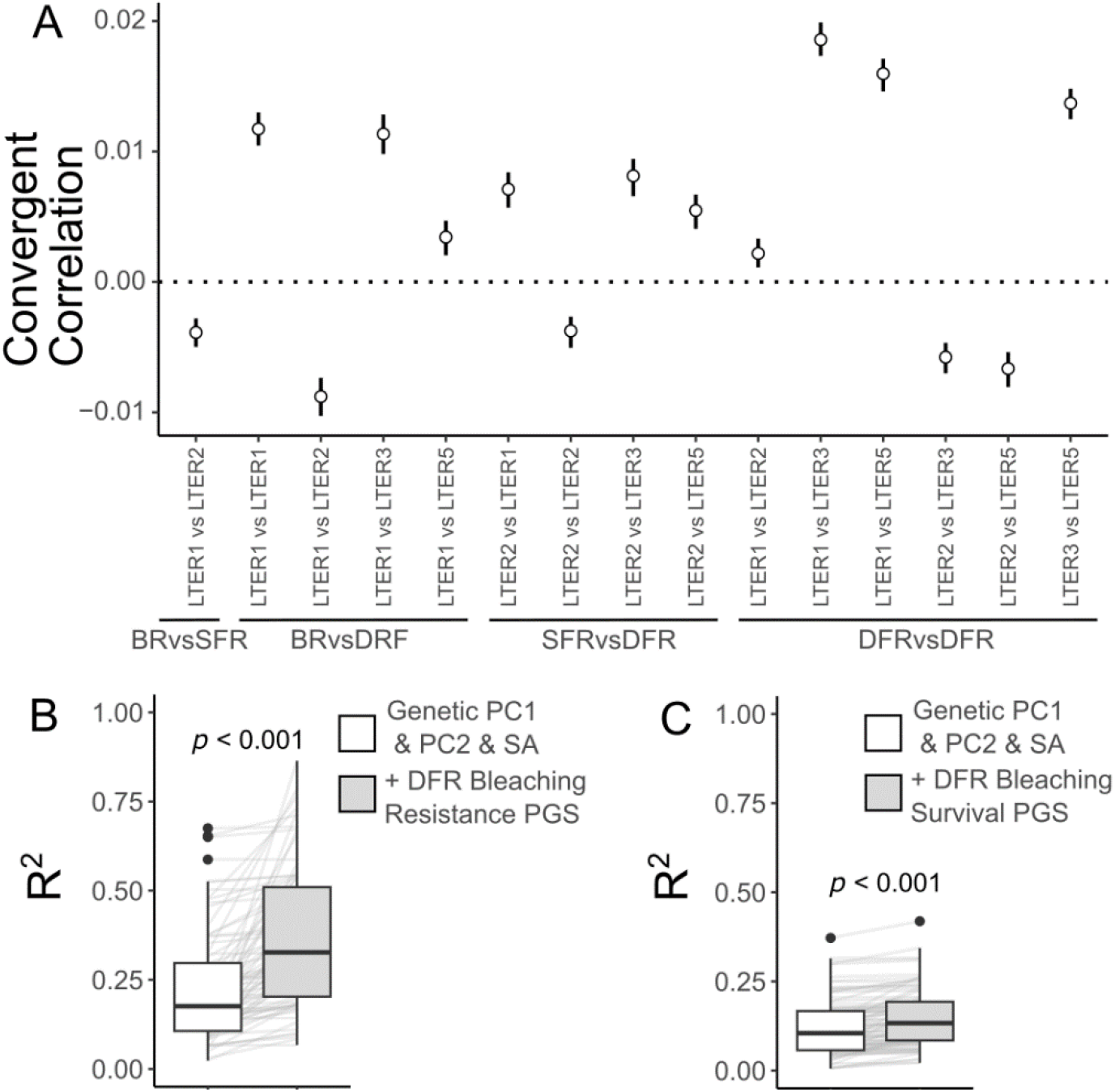
Bleaching resistance and survival have a host genetic basis. A) Convergent correlation statistic for allele frequency shifts between pre- and post-mortality timepoints. Vertical lines are 97.5% confidence intervals from 100 bootstraps. B) R^2^ distributions for test sets samples (100 reps, represented by horizontal grey lines) comparing each PGS (white) to randomly selected loci (light grey) for the bleaching resistance PGS and C) bleaching survival PGS. BR: Backreef; SFR: Shallow forereef; DFR: Deep forereef. SA: Surface Area.

We also calculated four PGSs (bleaching resistance PGS, bleaching survival PGS, deep forereef bleaching resistance PGS and deep forereef bleaching survival PGS). We show on its own the PGS offers very little predictability (mean R^2^s: 0.01-0.04; Supp. Fig. 15) and only the deep forereef PGSs offered significant increases in predictability compared to randomly selected loci (Supp. Fig. 16). However, given that both environmental and genetic factors contribute to variation in bleaching, it is important to determine whether the PGS increases predictability when accounting for other non-genetic factors. To do so we combined our PGS with the same variables included as covariates, to assess whether including these parameters improved model prediction. We also ensure that the improvement in model prediction offered by the PGS is not due to overfitting by comparing the model’s performance when we substitute the real PGS with a null PGS (*i.e.* randomly selected loci). For deep forereef PGSs only, we found that adding the respective PGS (bleaching resistance or bleaching survival) to a model with genetic PCs, colony size, and depth significantly improved the prediction of both health score (p < 0.05) and bleaching survival (p < 0.05), with this increase in predictive ability being greater than that offered by null PGSs (Supp. Fig. 17; Fig. 3B; Fig. 3C).

We also found that adding the respective PGS to a model with genetic PCs, colony size, and depth as well as both *Symbiodinium* and *Cladocopium* proportions, increases the predictability for the deep forereef bleaching resistance PGS but not for the other PGSs (Supp. Fig. 18; Supp. Fig. 19). However, it is important to note that pre-mortality symbiont communities are not necessarily indicative of what the coral hosted before the heat stress event. In fact, the strong correlation between having a mixed symbiont community and showing a bleached phenotype (Fig. 5A), suggests the predictive power of symbiont genera here is heavily inflated and thus we would not necessarily expect the PGS to add predictive power for this comparison.

### Reef-habitat-specific responses to the bleaching event

While there is some degree of convergence between the deep forereef sites and the backreef and shallow forereef in their response to the MME (Fig. 3A), we also find evidence that hosts from different reef habitats had unique responses. First, as described above, the deep forereef PGSs were able to offer predictability while the PGSs generated from all reef habitats were not (Supp. Fig. 17). Second, we observe a lack of convergent evolution between backreef and shallow forereef populations both genome-wide (Fig. 3A) and at the loci for the deep forereef bleaching survival PGS (Supp. Fig. 20). Third, a deep forereef bleaching survival PGS (*i.e.* a PGS generated for deep forereef individuals only with survival as the trait) appears to offer no increase in predictive power for determining survivorship in backreef individuals compared to PGSs built from randomly selected loci (Supp. Fig. 21). Fourth, we only find putative major effect loci at the backreef. It is possible that the reef-habitat specificity is due to opposing selection pressures, which aligns with our findings from the symbiont populations suggesting some degree of local adaptation between environments. In other words, a heat stress event might not impose a single homogenous selective pressure across all corals from impacted habitats. For example, during the heat stress event the shallow forereef experiences higher irradiance in comparison to the deeper forereef habitat (Dubé et al., 2023), an abiotic factor that impacts Symbiodiniaceae physiology by different mechanisms compared to heat stress (Downs et al., 2013). These diverse environments likely result in variable selective pressures across habitats, each imposing unique influences on the coral genome. It is also possible that there is a high degree of genetic redundancy for bleaching resistance and survival and reef-habitat specific shifts are due to differences in standing genetic variation. This theory is supported by the lack of a convergent shift in allele frequencies between backreef and shallow forereef habitats, which also happens to be the comparison with the greatest degree of genetic structure (Supp. Fig. 3).

### Signatures of selection and identification of adaptive variants in the next generation following a MME

To test whether allele frequency shifts resulting from the MME are maintained in subsequent generations, we examined the effect sizes of a pre-mortality (deep forereef only) vs juvenile GWAS at the roughly 150,000 loci from the mortality deep forereef PGS in Figure 3C. We find the sum of effect sizes at these loci was higher than randomly selected loci (Fig. 4A), demonstrating these loci were also important for explaining pre-mortality vs juvenile genetic differentiation. We then used DAPC analyses for the ∼150,000 loci in the bleaching survival PGS to investigate whether juveniles maintain the degree of divergence at these loci or if allele frequencies return towards pre-mortality levels. DAPC results using pre-mortality and post-mortality samples as a training dataset show that juveniles represented an intermediate point between the two timepoints on LD1 (Fig. 4B; Supp. Fig. 22). When naively assigning juveniles to either pre- or post-mortality groupings, 32/74 juveniles assigned to the pre-mortality group and 42/74 assigned to the post-mortality group (Supp. Fig. 22).

**Figure 4.**
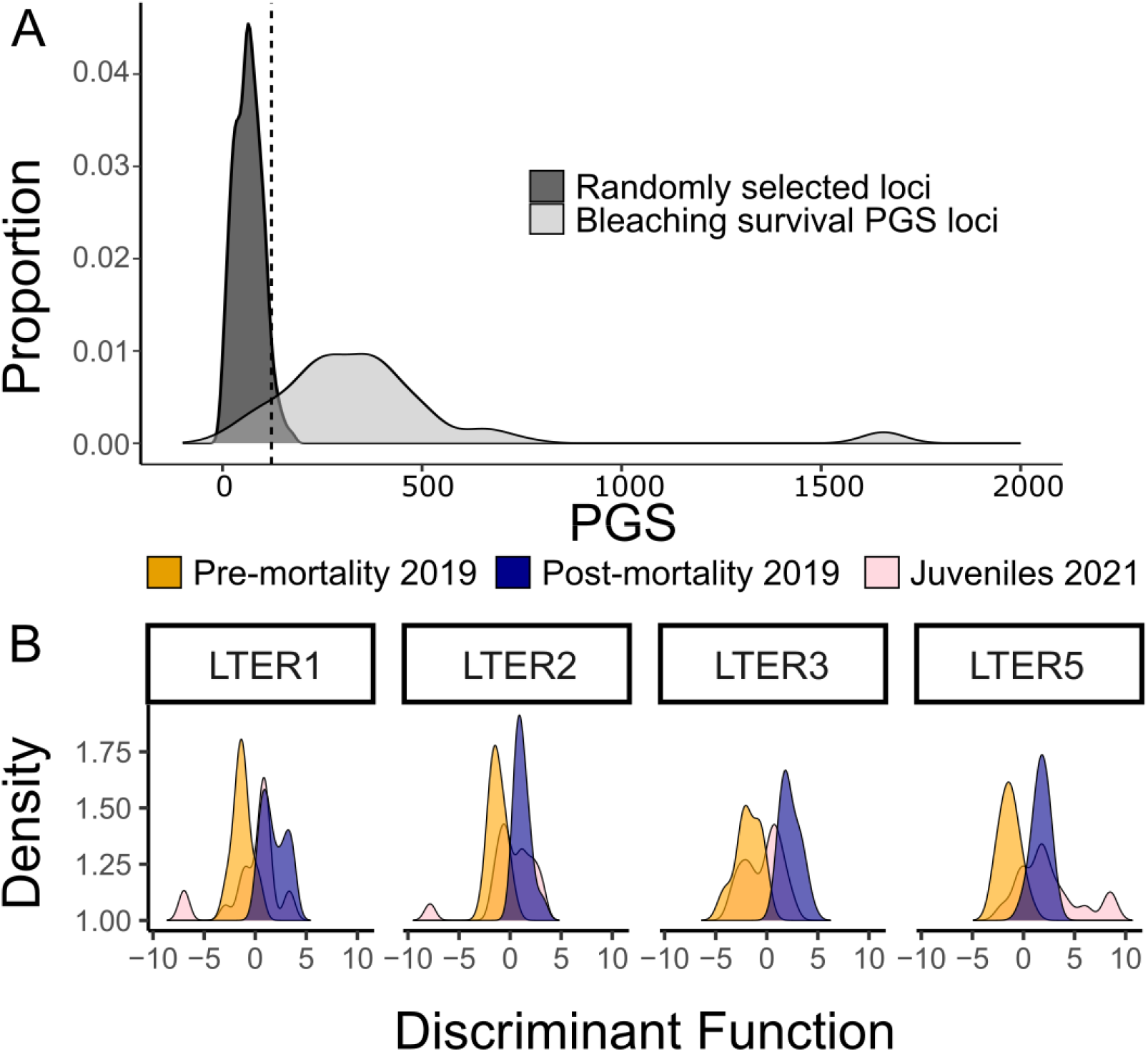
Juveniles maintain divergence in loci that predict bleaching survival. A) Distribution of summed beta scores from pre-mortality vs juveniles GWAS for both 500 sets of randomly selected loci (dark grey) and the loci used for the deep forereef bleaching survival PGS (Fig. 3C; light grey). Vertical line shows 95^th^ percentile for the randomly selected loci. B) DAPC results for the loci significantly differentiated between timepoints per GWAS, demonstrating the intermediate status of juvenile samples.

One possible interpretation of this result is that the allele frequency shifts caused by the MME are mitigated in the following generation, either through genetic drift or an opposing selective pressure (*e.g.*, competition for space). Alternatively, it is possible that this “intermediate frequency” in the following generation is actually the expected outcome under a scenario of rapid adaptive evolution if we consider the quantitative genetics model used here to capture “outlier loci” between pre- and post-mortality timepoints will contain false positives due to sampling bias from not capturing the entire population. We show indeed that we can recreate this intermediate frequency in outlier loci in the generation after an MME by simulating adaptive evolution in a population under our sampling scheme (Supp. Fig. 23).

Another possible interpretation is that these putatively adaptive loci are only shifting due to the high recruitment from the location (LTER 3) that experienced the least heat stress (Fig. 2D). In other words, frequency shifts are driven by individuals who were not simply exposed to adequate heat stress rather than those who were exposed and survived. We do not believe this is the case for two major reasons: 1) These adaptive genetic loci exhibit a convergent shift between all deep forereef sites (Supp. Fig 20), which we would not expect if shifts were due to genetic drift (Buffalo & Coop, 2020) and 2) LTER 3 at the pre-mortality timepoint should also show the intermediate frequency observed in juveniles if shifts are simply due to recruitment from LTER 3, but this pattern was not observed (Supp. Fig. 24).

Our simulations suggest that the loci exhibiting divergence in the juveniles compared to the pre-mortality timepoint may provide a better representation of the true adaptive loci for this MME. Given that GWAS effect sizes are obtained as a noisy combination of the unobserved true (single-nucleotide polymorphism) SNP effect sizes (Hormozdiari et al., 2014; Zhu et al., 2018), identifying true adaptive loci by taking the overlap of a GWAS between pre- and post-mortality and a second GWAS between pre-mortality and juveniles (both using deep forereef sites only) is likely overly conservative. However, our approach will still capture putatively important loci for adaptation, with the caveat that it may lead to false negatives. 316 loci passed the 0.01 *p*-value threshold for both the deep forereef bleaching survival GWAS and the pre-mortality vs juvenile GWAS (Supp. File 1). It is important to note that none of these loci project to be of major effect (as outlined previously in “Host shows highly polygenic response to the bleaching event”). Of the overlapping loci with the highest signal (*i.e.*, *p* <0.001) in the pre-mortality vs juvenile GWAS, we found several loci with previously described functions in cnidarians including Transcription factor AP2 (TFAP2A), Notch receptor 2 (NOTCH2), and NF-kappa-B inhibitor-interacting Ras 2 (NKIRAS2). TFAP2A is essential for germ cell commitment and gonad development in cnidarians (DuBuc et al 2020). NOTCH2 is important during development for cell fate determination, neurogenesis, and for establishing tissue boundaries during the budding of new polyps in cnidarians (Käsbauer et al., 2007; Marlow et al., 2012; Münder et al., 2010). NKIRAS2 inhibits NFkB activation (Tago et al., 2010) which is a transcription factor involved in stress response and innate immunity across the metazoan tree of life (Gilmore & Wolenski, 2012; Weis, 2019). These genes are causal candidates and could prove to be useful for future functional research on coral bleaching.

### Local adaptation of *Symbiodinium and Cladocopium* between shallow and deep forereef sites

Reads mapping to the genomes of three Symbiodiniaceae genera (*Symbiodinium*, *Cladocopium*, and *Durusdinium*) were used to approximate the proportion of symbionts in each sample. It is important to note that our “pre-mortality” timepoint occurs during the bleaching event, and therefore does not reflect the symbiont community prior to heat stress. This has two major implications for interpreting our results. 1) It is unclear whether the environmental structuring of symbiont communities described below is observed year-round or is specific to the heat stress event (see Leinbach et al., 2023 for a detailed discussion). 2) Bleached corals across habitats generally exhibited more mixed communities compared to healthy corals (Fig. 5A). However, the cause-effect relationship for the bleached corals (*i.e.*, low-health corals) is not clear—in other words, it is uncertain whether the observed symbiont community structure caused the bleaching or is a symptom of dysbiosis. Despite this uncertainty, bleached corals still provide valuable insights into the presence of specific symbiont genera at each reef habitat. Pairing this information alongside symbiont communities sampled from healthy corals during the bleaching event and from surviving corals post-bleaching allows us to investigate whether a particular symbiont genus might be locally adaptive.

**Figure 5.**
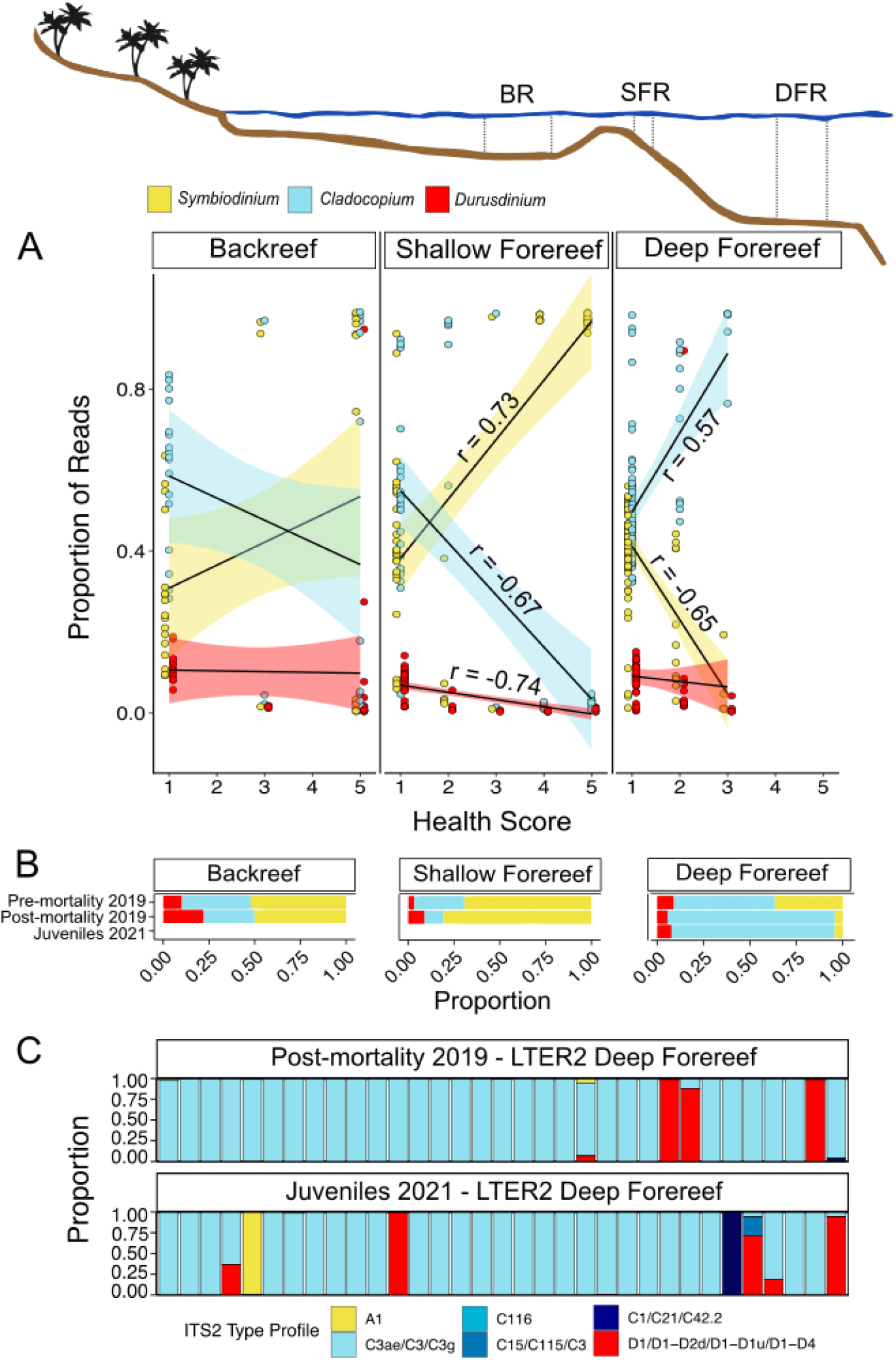
Algal symbiont divergence between reef zones contributes to bleaching response. A) Relationships between health score (1 indicates complete bleaching and 5 indicates no pigmentation loss (Photo examples in Supp. Fig. 12)) and Symbiodiniaceae genera during the bleaching event (*i.e.* pre-mortality May 2019) at backreef, shallow forereef and deep forereef habitats. Each dot represents the relative proportion of mapped reads to the three Symbiodiniaceae genera. Black lines show fitted linear regression with 95% confidence intervals. r values show Pearson correlation coefficient and only significant regressions are shown. B) Proportion of reads mapped to each of the three Symbiodiniaceae genera (*Symbiodinium, Cladocopium,* and *Durusdinium*) for each site across timepoints at backreef, shallow forereef, and deep forereef habitats. C) Proportion of collapsed ITS2 type profiles for individual colonies at LTER 2 deep forereef for both 2019 post-mortality (Leinbach et al., 2023) and 2021 timepoints.

We find evidence for locally adaptive symbiont genera between the shallow and deep forereefs. At shallow forereef sites (3-5 m), increased *Symbiodinium* reads were associated with lower bleaching susceptibility (*i.e.*, higher health score; *p* < 0.0001; r = 0.73), whereas increased *Cladocopium* reads were associated with higher bleaching susceptibility (*i.e.*, lower health score; *p* < 0.0001; r = −0.67; Fig. 5A). These results are consistent with characterizations of symbiont communities using ITS2 amplicon sequencing from recovered and resistant colonies at the LTER 2 site (Leinbach et al., 2023). However, the deep forereef showed the opposite trend, where an increased abundance of *Cladocopium* reads were associated with lower bleaching susceptibility (*p* < 0.0001; r = 0.57), while an increased abundance of *Symbiodinium* reads was associated with higher bleaching susceptibility (*p* < 0.0001; r = −0.65; Fig. 5A). The deep forereef showed changes in the proportion of Symbiodiniaceae genera between timepoints, with an increase in the proportion of *Cladocopium* post-mortality (October 2019) compared to pre-mortality (May 2019), and a corresponding decrease of both *Symbiodinium* and *Durusdinium* (*p* < 0.0001, Fig. 5B). It is possible symbiont zonation could be related to host genetic structuring (Bongaerts et al., 2010; Brazeau et al., 2013), however this reef system has high gene flow between habitats and little population structure, as shown in this study (Fig. 2A) and in Kriefall et al. (2022) so this explanation seems unlikely. This is further supported by results from a redundancy analysis (RDA) where host genetic structure was not explained by symbiont dominance (*Symbiodinium: p* = 0.614, R^2^ < 0.01; *Cladocopium*: *p* = 0.638, R^2^ < 0.01; *Durusdinium*: *p* = 0.434, R^2^ < 0.01; Supp. Table 2), pointing to a lack of host-driven symbiont structuring.

### Maintenance of symbiont community structure two years after the heat stress event

The increase of *Cladocopium* dominance on the deep forereef was maintained in 2019 juveniles, which exhibited no significant changes in *Cladocopium* from the post-bleaching October 2019 timepoint (*p* = 0.09, Fig. 5B). As there has never been information collected on the symbiont communities of *A. hyacinthus* from the deep forereef prior to the bleaching event in May 2019, we do not know if this structure maintains a post-bleaching shift or reflects historical structure. ITS2 amplicon sequencing of the collected juveniles in this study shows that *A. hyacinthus* at all deep forereef sites are dominated by *Cladocopium* C3ae, with occasional exceptions of certain colonies with multiple symbiont associations and a few dominated by A1, C1, and D1 (Fig. 5C; Supp. Fig. 25). We observe a negative correlation between juvenile colony size and ITS2 type richness (*p* < 0.05; Supp. Fig. 26), indicative of symbiont winnowing (Abrego et al., 2009), however this doesn’t appear to significantly increase symbiont diversity at the juvenile timepoint as the vast majority of juveniles sampled (96/115, 83.48%) were dominated by C3ae and we observed no differences in symbiont community composition between timepoints (*p* = 0.825, Supp. Fig. 27). *Cladocopium* C3ae is not prevalent in *A. hyacinthus* outside of Mo’orea, with the C3ae variant only appearing once in the Cook Islands (Lewis et al., 2022). However, without more comprehensive sampling at other islands, it is difficult to know whether the C3ae variant here is truly endemic to Mo’orea.

### No evidence for adaptive rescue by *Durusdinium*

Despite the presence of *Durusdinium* in corals from all reef habitats, we find no evidence that corals escaped or recovered from bleaching by shifting to *Durusdinium* dominance. This result contrasts the large body of observations that *Durusdinium* (formerly *Symbiodinium* Clade D) is associated with higher thermal tolerance, with a trade-off of lower growth rates, across diverse coral genera (Berkelmans & Van Oppen, 2006; Fuller et al., 2020; Klueter et al., 2017; LaJeunesse et al., 2018; Morikawa & Palumbi, 2019). In our study, *Durusdinium* was only correlated with health score at the shallow forereef, where a higher proportion was associated with more bleaching (*p* < 0.05, r = −0.74; Fig. 5A). It is unlikely that the *Durusdinium* we observe in *A. hyacinthus* is a novel, heat-susceptible species of *Durusdinium,* as ITS2 symbiont profiles for juvenile coral samples (Fig. 5C) and a subset of adult samples (Leinbach et al., 2023) contain both the D1 and D4 sequences characteristic of *D. trenchii* (Hume et al., 2020), which has previously been linked to bleaching resistance (Silverstein et al., 2017). Rather, it seems that we have found another rare exception to the rule of *Durusdinium* conferring holobiont heat tolerance (Abrego et al., 2008; Howe-Kerr et al., 2020; Howells et al., 2020). As stated previously, because data were generated from some severely bleached colonies, it is possible that the symbiont structure we observed pre-mortality (May 2019) is not indicative of the structure during or prior to the heat stress, but rather reflects symbionts that remained or were taken up once the colony bleached. However, if hosting *Durusdinium* was adaptive here, we would expect *Durusdinium* to increase in relative abundance from pre- to post-mortality timepoints, which we did not observe. In fact, at the deep forereef, we record a decrease in *Durusdinium* from pre- to post-mortality timepoints (*p* < 0.0001, Fig. 5B).

## DISCUSSION

Since the 1970s, coral reef populations in Mo’orea have experienced multiple cycles of disturbance and recovery (Adjeroud et al., 2009; Holbrook et al., 2018). These disturbance regimes, which include outbreaks of the corallivorous seastar *Acanthaster planci* in 1980-1982 (Berumen & Pratchett, 2006) and 2006-2009 (Kayal et al., 2012), cyclones in 1983 (Harmelin-Vivien & Laboute, 1986), 1991 (Adjeroud et al., 2002), and 2010 (Kayal et al., 2012), and several heat stress events (Hoegh-Guldberg et al., 2007; Penin et al., 2007), were then followed by the return of coral cover to pre-disturbance levels within 8-10 years (Adjeroud et al., 2009; Holbrook et al., 2018). In other locations in the South Pacific, recovery of *A. hyacinthus* is thought to depend on a particularly heat resilient genetic lineage within the *A. hyacinthus* complex (Rose et al 2021), or availability of the most heat resilient symbionts (*e.g.*, *Durusdinium* (Oliver & Palumbi, 2011). The Mo’orean *A. hyacinthus* populations allow us to address an important question facing coral reefs in the Anthropocene: how will coral populations rapidly adapt to disturbances when they have already exhausted sources of adaptive variation from within the species complex (*i.e.*, this *A. hyacinthus* population only consists of corals from the “heat resilient” HE *A. hyacinthus* genetic lineage (Rose et al., 2021) and from the symbiont (*i.e.*, *Durusdinium* is present but fails to impact survival)? We find that survival is predicted through the small effect of multiple alleles that are maintained in the following generation, providing evidence for contemporary adaptation to rising temperatures.

Our data show there is a polygenic basis for bleaching in the host, in line with previous work (Dixon et al., 2015; Drury et al., 2022; Fuller et al., 2020; Kirk et al., 2018; Rose et al., 2018, 2021), as well as for survival from the bleaching event and that the genetic variation that predicts bleaching survival is maintained in juveniles two years later. While the contribution to bleaching predictability might be small, the finding that intraspecific host genetic variation impacts variation in mortality at all is significant. It is important to note that parameters such as size, depth and symbiont genera are relatively static compared to host genetic variation. If a population’s survival is limited to protection offered by non-genetic factors, this imposes a lower ceiling on the ability to persist in the face of continuous stressors and coral populations are not predicted to survive without adaptation driven by the host (Matz et al., 2020). This rapid evolution in response to the 2019 heat stress event may help explain why there was no further decline in *Acropora* coral cover following the 2020 heat stress event.

Further, this MME did not result in decreased genetic diversity in the short-term, likely due to large N_e_ sizes of *A. hyacinthus*. The stability of genetic diversity, in combination with robust recruitment (Leichter et al., 2013), could explain the rapid returns to baseline coral cover following disturbances consistently observed in Mo’orea. However, it remains to be seen if genetic diversity will be maintained even after high mortality in subsequent generations that experience increased frequency of disturbance events. Stability of genetic diversity following large disturbance has been observed in *A. spicifera* due to spatial variability of disturbance impact (Thomas et al., 2024), which we also see here to some extent. We find the sites with the highest mortality show lower genetic diversity in 2021 juveniles, while the site that experienced the lowest heat stress, bleaching, and mortality (LTER 3) served as the largest source for juvenile recruitment and had similar levels of genetic diversity in 2021 juveniles compared to pre- and post-mortality adult populations. This suggests that MMEs can alter recruitment dynamics simply through spatial variation in stressor severity.

Despite the encouraging evidence of a population evolving rather than perishing in the face of MHWs, there are two points that should temper expectations. First, it is important to note that even if adaptive responses occur in response to MMEs, population extinction or depletion to the point of extinction debt may still result. As environmental changes continue and accelerate, mean population phenotypes can increasingly lag behind the optimal phenotype, leading to an increasing burden of selective deaths (Hansen et al., 2012). Second, while a PGS has previously been proposed to provide a means to develop predictive models of bleaching for distinguishing tolerant individuals across different shelf positions, latitudes and environmental conditions (Fuller et al., 2020), we find evidence for localized selective pressures having disparate impacts on allele frequency shifts. The habitat-specific accuracy of our PGS observed here demonstrates the difficulty in applying a PGS across diverse datasets and populations. We also note that host genetic variation offers relatively small increases in predictability for bleaching and survival compared to symbiont associations and other non-genetic factors. This is not to say that this approach cannot lead to a robust predictive model, but rather stresses the need to first thoroughly sample across habitats and populations and models should not substitute genetic variation for important non-genetic factors when attempting to optimize bleaching predictions.

We confirm previous findings of *A. hyacinthus* bleaching resistance mediated through hosting environment-specific symbiont types (Leinbach et al., 2023) and further document a complete role reversal of resilient symbiont partners between shallow forereef and deep forereef sites. This stark difference in environment-specific responses to a heat stress event between different algal partners in a panmictic host population suggests symbionts are adaptive in one environment and maladaptive in the other. It is possible that this local specialization is driven in part by the differences in light intensity between habitats (Dubé et al., 2021), as this depth-related relationship between *Symbiodinium* and *Cladocopium* has previously been shown (Varasteh et al., 2022). The benefit of *Symbiodinium* at the shallower forereef might be due to their ability to efficiently synthesize photoprotectors as mycosporine-like amino acids (MAAs; (Silva-Lima et al., 2015)) and produce UV-protective MAAs and low amounts of hydrogen peroxide, a putative agent of coral bleaching, at elevated temperatures (Banaszak et al., 2000). While symbiont tradeoffs are often framed as one symbiont species being beneficial under environmental stress and another more beneficial when the stress has dissipated (*e.g.*, Jones & Berkelmans, 2010), here we show trade-offs do not have to be limited in this way. Selection on these symbiont-environment associations can occur in opposing directions during the same heat stress event in different reef habitats.

This temporal genomic approach reveals key insights into resilience mechanisms within coral populations amidst MMEs. We show evidence for discrete adaptive roles of symbiont types in distinct reef habitats, and the interplay between host and symbiont genetics, highlighting the complexity of coral survival strategies. Our PGS and selection scan approach provides novel insight into the genetic architecture of MME survival and demonstrates that selective pressures vary both spatially (*i.e.*, across habitats) and temporally (*i.e.*, during stressor and recovery phases of the MME). Lastly, we document adaptation to warming temperatures from standing within-species genetic variation. These findings highlight the intricate nature of selective pressures of MMEs, offering crucial insights into coral resilience and adaptation under ongoing, unprecedented warming.

## METHODS

### Coral collections

Samples were collected via SCUBA from *A. hyacinthus* colonies surrounding the island of Mo’orea as follows: 1) adults during the May 2019 bleaching event (pre-mortality; N = 172), 2) adults after bleaching and thermal stress had subsided in October 2019 (post-mortality; N = 103), and 3) juveniles in November 2021 (N = 115). Sampling in May was carried out to maximally capture the entire bleaching phenotypic landscape, with an effort to sample bleached and unbleached colonies at sites whenever possible. However, bleaching varied across sites, so it was not possible to sample equal numbers of corals across the bleaching spectrum. At the time of sampling in May, we did not observe the onset of mortality in any of the corals, despite many in a severely bleached state. Therefore, we strongly suspect we sampled prior to the onset of any mortality. Ten colonies were repeatedly sampled at both May 2019 and October 2019 timepoints.

At each LTER site, colonies were sampled from at least one of three reef habitats (backreef, shallow forereef, and deep forereef), corresponding to three different depths (1-3 m, 3-5 m, 10-14 m, respectively). While adults from May 2019 and October 2019 included sampling at all three reef habitats, juveniles from November 2021 were only sampled at the deep forereef (Supp. Table 1). Juveniles were characterized as small colonies < 8 cm long diameter, and care was taken to ensure they were not remnants of surviving adults. Most juveniles were observed and sampled on recently deceased *Pocillopora spp.* skeletons or coral rubble. Deceased *A. hyacinthus* skeletons are easily differentiated in this environment by their broad tabular morphology and any juvenile or live coral sample found on *A. hyacinthus* skeletons was not sampled to avoid potentially sampling surviving colony fragments. Photographs of sampled juveniles are available in data repository linked at https://github.com/jamesfifer/MooreaWGS. Each sample consisted of a 2-3 cm fragment, collected with bone cutters, immediately preserved in 200 proof ethanol, and stored at −80°C. For each colony sampled, photographs containing size and color standards were taken and colony area and diameters of short and long sides were calculated using ImageJ (Schneider et al., 2012). Health score was estimated visually by using a Coral Color Reference Card (Siebeck et al., 2006). Representative colonies for each health score are presented in Supplementary Figure 12. To better represent the impacts of thermal stress on bleaching severity, additional colonies (N = 226) were photographed only (*i.e.*, not sampled for sequencing) and assigned a health score (Supp. Table 1).

### Coral mortality and bleaching surveys

Coral bleaching and mortality surveys were conducted at 10 m on the forereef of Mo’orea between July 9-15, 2019. Divers on SCUBA conducted two 50-m transects at each site, in which they quantified bleaching and mortality for all *A. hyacinthus* along a 1-m belt along the transect. Divers assessed bleaching and mortality for all *A. hyacinthus* greater than 5 cm in diameter. For each colony, divers estimated the percentage of each individual that was bleached or recently dead (*i.e.* if a colony had 25% mortality and 25% bleaching of the colony area then that colony would have 50% bleaching and mortality). Portions of colonies were classified as “recently dead” if they had been colonized by turf algae but not yet colonized by macroalgae. We summed the percentage of each colony that was bleached or recently dead.

### Temperature data

Water temperature data were collected as part of the MCR LTER core time series data collection (Leichter et al., 2019). At four sites (Fig. 1A), a bottom-mounted thermistor attached at 2 m (backreef) and 10 m (forereef) depth recorded water temperatures every 20 minutes. Cumulative heat stress was calculated as the 12-week running sum of weekly average temperatures exceeding 29°C, a noted bleaching threshold for corals in Mo’orea (Pratchett et al., 2013), from November 1, 2018 to December 1, 2021. Because LTERs 1, 2, and 3 did not record thermistor data for the entire desired timeframe, LTER 0 was substituted for temperature data at LTERs 1 and 2, and LTER 4 was substituted for LTER 3. As LTER 0 is only a few hundred meters away from LTER 1 and our sampling location fell between the two thermistor locations, we use the label “LTER 1” to refer to temperature data that came from the thermistor at LTER 0 in an effort to simplify figures. Correlations during time intervals with overlapping available data between sites suggest temperatures at substituted sites were extremely similar (Supp. Fig. 28). To address the concern that similarities between thermistor readings may decrease during heat stress events, we used available data from the subsequent 2020 heat stress event to show that cumulative heat stress was not different between substituted sites (Supp. Fig. 29).

### Whole Genome Sequencing (WGS)

DNA was isolated using a modified phenol-chloroform extraction method (Davies et al., 2013) and then cleaned with DNA Clean and Concentrator kits (Zymo, Irvine, CA). Cleaned genomic DNA was sent to the University of California Davis Genome Center for library preparations using pooled Super-High-Throughput Shotgun sequencing for shallow WGS (mean 6x coverage; Supp. File 1). Libraries were sequenced on Illumina NovaSeq 6000 S4 flow cells using two lanes of paired-end 150-bp and one lane using paired-end 100-bp. Adapters were trimmed and reads were filtered using *fastp* (Chen et al., 2018) (Phred scores >= Q 20 and 40% unqualified reads threshold). Contamination from symbiotic DNA was filtered out by mapping trimmed reads to a concatenated genome of four Symbiodiniaceae genera: *Symbiodinium* (Aranda et al., 2016) *Breviolum* (Shoguchi et al., 2013), *Cladocopium* (H. Liu et al., 2018), and *Durusdinium* (Dougan et al., 2022) via bowtie2’s (Langmead & Salzberg L., 2013) --un-conc option. Symbiont-free paired-end reads were mapped to the *A. hyacinthus* genome (López-Nandam et al., 2023) using bowtie2 (parameters -I 0 -X 1500 --no-unal –fr) resulting in an average mapping rate of 88.4% to the coral genome (Supp. File 1). Aligned reads were then merged across the three lanes using samtools (Li et al., 2009). The *clipOverlap* function of bamUtil (Jun et al., 2015) clipped overlapping read pairs and PCR duplicates were removed using *MarkDuplicates* in Picard (Picard Toolkit, 2019).

Genotyping and identification of SNPs were performed using ANGSD (Korneliussen et al., 2014). Standard filtering that was used across all analyses unless otherwise specified included loci present in at least 80% of individuals, minimum mapping quality score of 20, minimum quality score of 25, strand bias *p*-value > 0.05, heterozygosity bias > 0.05, minor allele frequency (MAF) > 0.05, a SNP *p*-value of 1×10^-5^, removing all triallelic sites, removing reads having multiple best hits and lumped paralogs filter. Running these filters across all samples produced 5,872,614 SNPs.

### Clone identification and genetic structure between sites, habitats, and timepoints

Clones were detected using hierarchical clustering of samples based on pairwise identity by state (IBS) distances calculated in ANGSD. Technical replicates were used to identify appropriate height cutoffs for clone identification. To compare population structure between sites, habitats, and timepoints, our data were subset into the following datasets: 1) pre-mortality only (May 2019 samples), 2) post-mortality only (October 2019 samples), 3) juveniles only (November 2021 samples), 4) pre-mortality (May 2019) and post-mortality (October 2019) across only the common locations between the two timepoints (LTER 1 backreef and deep forereef, LTER 2 shallow forereef and deep forereef, LTER 3 deep forereef, and LTER 5 deep forereef), and 5) all three timepoints across the locations common between all timepoints (LTER 1 deep forereef, LTER 2 deep forereef, LTER 3 deep forereef, and LTER 5 deep forereef). For each of the five datasets, genetic structuring between sites (datasets 1-3) or timepoints (datasets 4 and 5) was compared using PCoA via a covariance matrix based on single-read resampling calculated in ANGSD. DAPC was also performed from the PCoA table. DAPCs were carried out using the *lda* function from the R (R Core Team, 2023) package MASS (Ripley et al., 2013), where the number of PCs to retain was determined using scree plots and CV (checking 1: N/3 PCs, where N is the total number of eigenvalues). Additionally, DAPCs were used to identify potential source populations for juveniles. To do so, DAPCs were performed using adult colonies from 2019 (both pre- and post-mortality) only as training data, with site as the classification variable. The resulting model was used to predict juvenile site assignment. Population structure was also assessed using NGSadmix (Skotte et al 2013). PCoAs incorporating WGS data from Rose et al. (2021) were used to confirm that all samples were *A. hyacinthus*.

Expected heterozygosity was calculated from each individual’s site frequency spectrum (SFS) (dividing singletons by all loci) generated by first calculating the site allele frequency (SAF) in ANGSD with no MAF filter and then the unfolded (using the *A. hyacinthus* reference genome (López-Nandam et al., 2023) as an ancestral reference) SFS in winsfs (Rasmussen et al., 2022). Differences in individual heterozygosities between sites and timepoints were calculated via Dunn’s test (1964) with a Benjamini-Hochberg multiple test correction. As this only assesses the increase or decrease of heterozygous individuals, we also looked at genetic diversity dynamics of the population by calculating nucleotide diversity (Watterson’s θ) for each site at each timepoint using ANGSD’s *thetastat* function on the SFS for each population (randomly downsized to the minimum number of individuals for each site across timepoints). A similar method was also used to calculate π, using the pairwise θ from ANGSD’s *thetastat* function as described in Adams et al., (2023). To examine genetic differentiation between sites, SFSs were used as priors with the SAF to calculate global *F*_ST_. Here, only weighted global *F*_ST_ values between populations are reported. To determine how large *A. hyacinthus* N_e_ are compared to other *A. hyacinthus* populations, StairwayPlot V2 (Liu & Fu, 2020) was used to model N_e_ through time. Unfolded SFSs for N_e_ estimates were generated by first identifying a set of putatively unlinked loci through linkage disequilibrium (LD) pruning in PLINK2 (window size 200kb, step size 20 and r^2^ threshold of 0.2) and using a per generation mutation rate of 2*e*-8 and a generation time of 5 (Fifer et al., 2022).

### Genome-wide association study (GWAS) on variation in bleaching resistance in pre-mortality samples and bleaching survival in pre- and post-mortality samples

A GWAS for bleaching resistance was performed for pre-mortality samples taken during the bleaching event (N = 172), with health score as the trait, serving as a proxy for bleaching resistance (Fig. 1C). General linear models (GLMs) were used to test for additive effects of SNPs (with minor allele frequencies > 5%) on the quantile-normalized health score (Zhou et al., 2017; Zhou & Stephens, 2012), including as covariates the first two genetic PCs, non-genetic/environmental variables (surface area of the colony and the collection depth), and the proportion of *Symbiodinium* and *Cladocopium* reads (as two separate terms) relative to all symbiont reads. Reads mapping to three Symbiodiniaceae genera genomes (*i.e.*, *Symbiodinium, Cladocopium,* and *Durusdinium*) were used as an approximation of the proportion of symbionts in each sample following Fuller et al. (2020). Additional filtering included removing sites with more than 10% individuals missing and filtering out SNPs with Hardy-Weinberg *p*-values below 1*e*-7. For the GLM, a GWAS was performed using a standard linear regression as implemented in PLINK2. Genome-wide cutoffs were determined through 10,000 permutations of the GLM, randomly shuffling the trait values, extracting the minimum *p*-value for each run, and then taking the value of the 95^th^ percentile of this distribution (Fuller et al., 2020).

A GWAS was also carried out assigning the pre- and post-mortality timepoints as a binary trait (*i.e.*, bleaching survival) to calculate a GWAS for bleaching survival. An alternative design here would have been to follow samples and score whether they survived or not, however it is not necessary to follow the same samples to detect selection in a population following a mortality event (*e.g.*, Schiebelhut et al., 2018). Additionally, following a set of samples is not necessarily less biased, as it would be biased towards that set of individuals instead of the larger population. Here we use a logistical regression model in PLINK to accommodate the binary trait and repeat all covariates listed above. For the bleaching survival GWAS we also omitted colonies from the pre-mortality timepoint that were also present at the post-mortality timepoint (10 colonies). We scored these colonies only as post-mortality given all colonies from the post-mortality timepoint were also technically present at the pre-mortality timepoint and marking these colonies in both timepoints would not accurately score them as survivors. We also exclude all LTER 2 colonies from the post-mortality timepoint as complete mortality at the original site at 10 m necessitated sampling from 5 m deeper (*i.e.* at 15 m). Lastly, we also repeated both the bleaching resistance and bleaching survival GWAS subsetting the data into shallow forereef, deep forereef or backreef only. Information on which samples were included in each GWAS is available in Supplemental File 1.

### Investigating convergent signature of selection following mortality event

We can leverage the temporal sampling at multiple sites to examine covariance in allele frequency shifts following the mortality event. If shifts are caused by a common selective pressure we expect to observe concordance between locations whereas discordance between locations is reflective of genetic drift, opposing selection pressures or differences in standing genetic variation (Buffalo & Coop, 2020). To assess evidence for parallel adaptation, we used the “convergent correlation” statistic described by Buffalo and Coop (2020). This takes allele frequency shifts between timepoints at each location throughout the entire genome and assesses correlation using a Pearson correlation test. We estimated 95% bootstrap confidence intervals by resampling the loci used to calculate each convergent correlation statistic 100 times with replacement following Reid et al. (2023).

### Calculation of bleaching resistance and bleaching survival PGSs

We calculated four PGSs in total: bleaching resistance PGS, bleaching survival PGS, deep forereef bleaching resistance PGS and deep forereef bleaching survival PGS. For the deep forereef PGSs we only use samples from the deep forereef. All PGSs were calculated using a jackknife CV procedure (Fuller et al., 2020). Samples were subset into 100 partitions of training and test sets, withholding 15% of individuals for the test and using the rest (85%) as the training set. For each training set, a GWAS (using either the GLM or LM as described above) was performed. *P*-value thresholding was used to build the PGS (*p*-value thresholds considered: 1*e*-1, 1*e*-2, 1*e*-3, 1*e*-4, 1*e*-5 and 1*e*-6), followed by LD-clumping via PLINK (with r^2^ threshold of 0.2) to find approximately independent SNPs. For each test set, the PGS was calculated by summing the allelic dosages weighted by the estimated effect size. PGS accuracy of different *p*-value thresholds was measured by taking R^2^ from a linear model run with PGS as the sole parameter, and then proceeding with the *p*-value that maximized R^2^ (Supp. Fig. 15). We further assess the performance of the PGS on its own by comparing the distribution of R^2^s in the 100 test sets with R^2^s from a null PGS calculated by randomly selecting loci (with the same *n loci* as the real PGS). We then test whether the real PGS adds significant predictability compared to the null PGS using Mann-Whitney U tests.

To follow up on habitat-specific differences we examined whether the deep forereef bleaching survival PGS could predict survival at the backreef. This involves taking beta values calculated from the deep forereef bleaching survival PGS and applying them in an additive model to generate a score for backreef samples. We then calculated the R^2^ using a linear model to see how well the PGS predicted survival at the backreef and then compared this R^2^ to the distribution of R^2^s from 500 sets of randomly selected loci (with the same *n loci* as the real PGS). We repeated this with several different sets of loci and associated beta values for different *p*-value thresholds (1*e*-1, 1*e*-2, 1*e*-3, 1*e*-4).

Given that bleaching resistance and survival are likely influenced by a combination of genetic and non-genetic factors (Fuller et al., 2020) we do not necessarily expect the PGS to increase predictability on its own. Instead, it is more meaningful to assess whether a model with important non-genetic predictors of bleaching is improved when adding the PGS. We do so by assessing the change in R^2^ using Mann-Whitney U tests between linear models with different combinations of predictors (1. Genetic PCs, 2. Genetic PCs + Non-genetic/Environmental variables, 3. Genetic PCs + Non-genetic/Environmental variables + PGS, 4. Genetic PCs + Non-genetic/Environmental variables + Symbiont proportions, 5. Genetic PCs + Non-genetic/Environmental variables + Symbiont proportions + PGS). Specifically, comparisons were made between the predictability of model 5 versus model 4 and model 3 versus model 2, with the expectation that the PGS would increase the predictability of health score and survival. In instances where model 5 or 3 increased predictability relative to model 4 or 2, we also compared whether the respective model increased predictability when replacing the real PGS with a null PGS generated from randomly selecting the same number of loci.

### Assessing the distribution of adaptive variants in the next generation following a mortality event

To test whether allele frequency shifts resulting from the MME are maintained in subsequent generations, we first examined the effect sizes of a pre-mortality (using deep forereef sites only) vs juvenile GWAS at the ∼150,000 loci (before LD clumping) from the deep forereef bleaching survival PGS described above. The effect sizes at these loci were compared to 500 sets of 150,000 randomly selected loci and a 95^th^ percentile cutoff was used to determine significant deviation from this null distribution. To determine if allele frequencies after selection were maintained in the following generation (2021 juveniles), we performed PCoAs using the set of high signal loci identified by the deep forereef bleaching survival PGS and dataset 5 (all three timepoints across the locations common between all timepoints). We also performed DAPCs on the PCoA table as described in the section *Clone identification and genetic structure between sites, habitats, and timepoints*. Additionally, for the DAPC analysis, juveniles were excluded from the training data and then assigned to either timepoint using the model to measure greater proximity to either pre- or post-mortality groups.

To determine the expected distribution of allele frequency shifts in high signal loci from the deep forereef bleaching survival GWAS between timepoints (and compare with DAPCs above), simulations were performed, iterating over several values of population size and false positive rates. To simulate genotypes, random numbers were drawn from a binominal distribution with a probability of 0.1 per alternative allele for both simulated neutral and adaptive loci. Individuals in the top 30% of polygenic scores (to reflect ∼30% survival rates; Fig. 1D) were then randomly selected (to reflect subsampling of the true population) and a logistic regression between the simulated pre- and post-mortality timepoints (as carried out in the GWAS) was conducted to identify putative outlier loci. These outlier loci contain true adaptive loci (*i.e.*, adaptive in the whole population) or false positives (identified as outliers due to subsampling of the whole population). For each individual, a binomial distribution was again used to randomly assign alleles from the surviving genotypes for the next generation. We then visualized the distribution of these loci across both pre-, post-mortality and juvenile timepoints.

To identify putatively adaptive loci that maintained post-mortality allele frequencies in the subsequent generation we carried out another GWAS, this time assigning the pre-mortality (but using only deep forereef sites) and juvenile timepoints as a binary trait and using the same covariates described in the GWASs outlined above. Loci that passed the *p*-value cutoff of 1*e*-2 here and also in the deep forereef bleaching survival GWAS were identified as “overlapping loci” and annotated genes were identified +/− 500kb from the overlapping loci.

### Role of Symbiodiniaceae in bleaching response

Pearson’s correlation test (Freedman, 2007) was used to examine the relationship between colony health score and proportion of symbiont genus (determined using mapped reads to Symbiodiniaceae genera genomes as described above). Mann-Whitney U tests validated changes in proportion of each Symbiodiniaceae genus between the three timepoints. Reads were mapped to only three Symbiodiniaceae genera genomes because of their consistent presence in ITS2 data, with a noticeable lack of any detectable *Breviolum* symbiont types (Leinbach et al., 2023, Supp. Fig. 25).

To identify if host genetic variation could be structuring symbiont communities, we performed a RDA from the symbiont dominance information gathered from WGS reads. RDA models were conducted with individual-based genotypes; we imputed the most common genotype at each SNP across all individuals for missing data. A genotype matrix (012 allele coding system) was created for each lineage as the response variable for RDAs. We used the host genotype matrix as the response variable and symbiont dominance (*i.e.*, which symbiont had the highest proportion of reads) as the explanatory variable; then significance was examined with an ANOVA.

Juveniles collected in 2021 were also sequenced for ITS2 (N = 115, Supp. Table 1). ITS2 amplicons were generated via PCR using Symbiodiniaceae-specific primers SYM_VAR_5.8S2 and SYM_VAR_REV (Hume et al., 2018). Libraries were sequenced on the Illumina MiSeq platform with 300-bp paired-end reads at the Georgia Genomics and Bioinformatics Core at the University of Georgia. Paired forward and reverse ITS2 sequences from juveniles combined with ITS2 sequences from post-mortality timepoints (August 2019 (N = 16) and October 2019 (N = 58); (Leinbach et al., 2023)) were submitted to SymPortal (Hume et al., 2019) to predict ITS2 type profiles, which represent putatively different taxa.

After combining the 2019 and 2021 juvenile ITS2 data, a total of 78 ITS2 type profiles were produced from SymPortal. ITS2 type profile reads were normalized using the trimmed mean of M-values (TMM) method, using function *calcNormFactors* in the edgeR package (Anders et al., 2010) to adjust for differences in sequencing depth. The 78 ITS2 type profiles were collapsed into nine distinct profiles based on PCoAs of their Bray-Curtis indices (Supp. Fig. 30), then used in subsequent analyses. One profile (F3x) was excluded from analyses due to low read counts in a single sample (N2F57J). To explore how timepoint impacted symbiont community structure at the site for which there was data at both timepoints (LTER 2 deep forereef; six distinct ITS2 profiles), a NMDS plot was generated. Bray-Curtis dissimilarities were calculated from a matrix of ITS2 profile abundance for each sample. Multivariate dispersion was measured using the function *betadisper* on Bray-Curtis dissimilarities and permutational analysis of variances (PERMANOVAs), using the function *adonis* in the vegan package (Okansen et al., 2020), to assess changes in symbiont community structure between post-mortality and juvenile timepoints. We employed a linear mixed-effects model (LME) to analyze the relationship between ITS2 type profile richness and colony size and a linear model (LM) to determine if colony size differed by site.

## DATA AVAILABILITY

All analysis pipelines are open source and can be found at https://github.com/jamesfifer/MooreaWGS. Raw read data will be made available on NCBI’s sequence read archive (SRA) upon publication.

## ACKNOWLEDGEMENTS

This project was supported by the U.S. National Science Foundation (NSF) (OCE #1935308 to MES and OCE #1935305 to GEH) and startup funds from Texas A&M University to MES. This manuscript also uses data collected by the NSF’s Moorea Coral Reef Long Term Ecological Research (MCR LTER) site under award OCE #1637396 (and earlier awards). Additional financial support to the MCR LTER site was provided through a generous gift from the Gordon and Betty Moore Foundation. This research was completed under permits issued by the French Polynesian Government (Délégation à la Recherche) and the Haut-Commissariat de la République en Polynésie Française (DTRT) (Protocole d’Accueil 2019, 2021). Samples were exported to the United States under CITES (No.FR1998700174-E). In addition to co-authors, DNA extractions were supplementally performed by Megan Maloney, Maddie Grace, Marie Harris, Avery von Eiff and Kimberly Candelario. We would also like to acknowledge Terrance Leach, Logan Kozal, and Jannine Chamorro for assistance with field work, and Zach Fuller, Mikhail Matz, the Davies Lab and the Rose Lab for their feedback on analyses and figures.

## REFERENCES

Abrego, D., Ulstrup, K. E., Willis, B. L., & Van Oppen, M. J. H. (2008). Species-specific interactions between algal endosymbionts and coral hosts define their bleaching response to heat and light stress. Proceedings of the Royal Society B: Biological Sciences, 275(1648), 2273–2282. 10.1098/rspb.2008.0180

Abrego, D., Van Oppen, M. J. H., & Willis, B. L. (2009). Onset of algal endosymbiont specificity varies among closely related species of Acropora corals during early ontogeny. Molecular Ecology, 18(16), 3532–3543. 10.1111/j.1365-294X.2009.04276.x

Adams, N. E., Bandivadekar, R. R., Battey, C. J., Clark, M. W., Epperly, K., Ruegg, K., Tell, L. A., & Bay, R. A. (2023). Widespread gene flow following range expansion in Anna’s Hummingbird. Molecular Ecology, 32(12), 3089–3101. 10.1111/mec.16928

Adjeroud, M., Augustin, D., Galzin, R., & Salvat, B. (2002). Natural disturbances and interannual variability of coral reef communities on the outer slope of Tiahura (Moorea, French Polynesia): 1991 to 1997. Marine Ecology Progress Series, 237, 121–131. 10.3354/meps237121

Adjeroud, M., Michonneau, F., Edmunds, P. J., Chancerelle, Y., de Loma, T. L., Penin, L., Thibaut, L., Vidal-Dupiol, J., Salvat, B., & Galzin, R. (2009). Recurrent disturbances, recovery trajectories, and resilience of coral assemblages on a South Central Pacific reef. In Coral Reefs (Vol. 28, Issue 3, pp. 775–780). Springer Verlag. 10.1007/s00338-009-0515-7

Anders, S., Huber, W., Nagalakshmi, U., Wang, Z., Waern, K., Shou, C., Raha, D., Gerstein, M., Snyder, M., Mortazavi, A., Williams, B., McCue, K., Schaeffer, L., Wold, B., Robertson, G., Hirst, M., Bainbridge, M., Bilenky, M., Zhao, Y., … Salzberg, S. (2010). Differential expression analysis for sequence count data. Genome Biology, 11(10), R106. 10.1186/gb-2010-11-10-r106

Aranda, M., Li, Y., Liew, Y. J., Baumgarten, S., Simakov, O., Wilson, M. C., Piel, J., Ashoor, H., Bougouffa, S., Bajic, V. B., Ryu, T., Ravasi, T., Bayer, T., Micklem, G., Kim, H., Bhak, J., LaJeunesse, T. C., & Voolstra, C. R. (2016). Genomes of coral dinoflagellate symbionts highlight evolutionary adaptations conducive to a symbiotic lifestyle. Scientific Reports, 6(August), 1–15. 10.1038/srep39734

Arnaud-Haond, S., Duarte, C. M., Teixeira, S., Massa, S. I., Terrados, J., Tri, N. H., Hong, P. N., & Serrão, E. A. (2009). Genetic recolonization of mangrove: Genetic diversity still increasing in the Mekong delta 30 years after Agent Orange. Marine Ecology Progress Series, 390, 129–135. 10.3354/meps08183

Auteri, G. G., & Knowles, L. L. (2020). Decimated little brown bats show potential for adaptive change. Scientific Reports, 10(1). 10.1038/s41598-020-59797-4

Bairos-Novak, K. R., Hoogenboom, M. O., van Oppen, M. J. H., & Connolly, S. R. (2021). Coral adaptation to climate change: Meta-analysis reveals high heritability across multiple traits. In Global Change Biology (Vol. 27, Issue 22, pp. 5694–5710). John Wiley and Sons Inc. 10.1111/gcb.15829

Baker, A. C., Starger, C. J., McClanahan, T. R., & Glynn, P. W. (2004). Corals’ adaptive response to climate change. Nature, 430(7001), 741–741. 10.1038/430741a

Banaszak, A. T., Lajeunesse, T. C., & Trench, R. K. (2000). The synthesis of mycosporine-like amino acids (MAAs) by cultured, symbiotic dinoflagellates. In Journal of Experimental Marine Biology and Ecology (Vol. 249). www.elsevier.nl/locate/jembe

Barfield, S., Davies, S. W., & Matz, M. V. (2022). Evidence of sweepstakes reproductive success in a broadcast spawning coral and its implications for coral metapopulation persistence. Molecular Ecology. 10.1111/mec.16774

Berkelmans, R., & Van Oppen, M. J. H. (2006). The role of zooxanthellae in the thermal tolerance of corals: A “nugget of hope” for coral reefs in an era of climate change. Proceedings of the Royal Society B: Biological Sciences, 273(1599), 2305–2312. 10.1098/rspb.2006.3567

Berumen, M. L., & Pratchett, M. S. (2006). Recovery without resilience: Persistent disturbance and long-term shifts in the structure of fish and coral communities at Tiahura Reef, Moorea. Coral Reefs, 25(4), 647–653. 10.1007/s00338-006-0145-2

Bongaerts, P., Riginos, C., Ridgway, T., Sampayo, E. M., Van, M. J. H., Englebert, N., Vermeulen, F., & Hoegh-guldberg, O. (2010). Genetic Divergence across Habitats in the Widespread Coral Seriatopora hystrix and Its Associated Symbiodinium. 5(5). 10.1371/journal.pone.0010871

Brazeau, D. A., Lesser, M. P., & Slattery, M. (2013). Genetic Structure in the Coral, Montastraea cavernosa: Assessing Genetic Differentiation among and within Mesophotic Reefs. PLoS ONE, 8(5). 10.1371/journal.pone.0065845

Buffalo, V., & Coop, G. (2020). Estimating the genome-wide contribution of selection to temporal allele frequency change. 117. 10.1073/pnas.1919039117/-/DCSupplemental.y

Campbell-Staton, S. C., Cheviron, Z. A., Rochette, N., Catchen, J., Losos, J. B., & Edwards, S. V. (2017). Winter storms drive rapid phenotypic, regulatory, and genomic shifts in the green anole lizard. Science, 357(6350), 495–498. 10.1126/science.aam5512

Chen, S., Zhou, Y., Chen, Y., & Gu, J. (2018). Fastp: An ultra-fast all-in-one FASTQ preprocessor. Bioinformatics, 34(17), i884–i890. 10.1093/bioinformatics/bty560

Coleman, M. A., Minne, A. J. P., Vranken, S., & Wernberg, T. (2020). Genetic tropicalisation following a marine heatwave. Scientific Reports, 10(1), 1–11. 10.1038/s41598-020-69665-w

Colson, I., & Hughes, R. N. (2004). Rapid recovery of genetic diversity of dogwhelk (Nucella lapillus L.) populations after local extinction and recolonization contradicts predictions from life-history characteristics. Molecular Ecology, 13(8), 2223–2233. 10.1111/j.1365-294X.2004.02245.x

Cooke, I., Ying, H., Forêt, S., Bongaerts, P., Strugnell, J. M., Simakov, O., Zhang, J., Field, M. A., Rodriguez-Lanetty, M., Bell, S. C., Bourne, D. G., Jh Van Oppen, M., Ragan, M. A., & Miller, D. J. (2020). Genomic signatures in the coral holobiont reveal host adaptations driven by Holocene climate change and reef specific symbionts. In Sci. Adv (Vol. 6). https://www.science.org

Davies, S. W., Rahman, M., Meyer, E., Green, E. A., Buschiazzo, E., Medina, M., & Matz, M. V. (2013). Novel polymorphic microsatellite markers for population genetics of the endangered Caribbean star coral, Montastraea faveolata. Marine Biodiversity, 43(2), 167–172. 10.1007/s12526-012-0133-4

Dixon, G. B., Davies, S. W., Aglyamova, G. V., Meyer, E., Bay, L. K., & Matz, M. V. (2015). Genomic determinants of coral heat tolerance across latitudes. Science, 348(6242), 1460– 1462. 10.1126/science.1261224

Dougan, K. E., González-Pech, R. A., Stephens, T. G., Shah, S., Chen, Y., Ragan, M. A., Bhattacharya, D., & Chan, C. X. (2022). Genome-powered classification of microbial eukaryotes: focus on coral algal symbionts. In Trends in Microbiology (Vol. 30, Issue 9, pp. 831–840). Elsevier Ltd. 10.1016/j.tim.2022.02.001

Douglas, A. E. (2003). Coral bleaching - How and why? In Marine Pollution Bulletin (Vol. 46, Issue 4, pp. 385–392). Elsevier Ltd. 10.1016/S0025-326X(03)00037-7

Downs, C. A., McDougall, K. E., Woodley, C. M., Fauth, J. E., Richmond, R. H., Kushmaro, A., Gibb, S. W., Loya, Y., Ostrander, G. K., & Kramarsky-Winter, E. (2013). Heat-stress and light-stress induce different cellular pathologies in the symbiotic dinoflagellate during coral bleaching. PLoS ONE, 8(12). 10.1371/journal.pone.0077173

Drury, C., Bean, N. K., Harris, C. I., Hancock, J. R., Huckeba, J., H, C. M., Roach, T. N. F., Quinn, R. A., & Gates, R. D. (2022). Intrapopulation adaptive variance supports thermal tolerance in a reef-building coral. Communications Biology, 5(1). 10.1038/s42003-022-03428-3

Du, L., Cai, C., Wu, S., Zhang, F., Hou, S., & Guo, W. (2016). Evaluation and exploration of favorable QTL alleles for salt stress related traits in cotton cultivars (G. hirsutum L). PLoS ONE, 11(3). 10.1371/journal.pone.0151076

Dubé, C. E., Hume, B. C., Boissin, E., Mercière, A., Bourmaud, A., Ziegler, M., & Voolstra, C. R. (2023). Algal symbioses with fire corals demonstrate host genotype specificity and niche adaptation at subspecies resolution. BioRxiv. 10.1101/2023.04.03.535406

Dubé, C. E., Ziegler, M., Mercière, A., Boissin, E., Planes, S., Bourmaud, C. A. F., & Voolstra, C. R. (2021). Naturally occurring fire coral clones demonstrate a genetic and environmental basis of microbiome composition. Nature Communications, 12(1). 10.1038/s41467-021-26543-x

Eakin, C. M., Sweatman, H. P. A., & Brainard, R. E. (2019). The 2014–2017 global-scale coral bleaching event: insights and impacts. Coral Reefs, 38(4), 539–545. 10.1007/s00338-019-01844-2

Fabricius, K. E. (2006). Effects of irradiance, flow, and colony pigmentation on the temperature microenvironment around corals: Implications for coral bleaching? Limnology and Oceanography, 51(1 I), 30–37. 10.4319/lo.2006.51.1.0030

Fey, S. B., Siepielski, A. M., Nusslé, S., Cervantes-Yoshida, K., Hwan, J. L., Huber, E. R., Fey, M. J., Catenazzi, A., & Carlson, S. M. (2015). Recent shifts in the occurrence, cause, and magnitude of animal mass mortality events. Proceedings of the National Academy of Sciences of the United States of America, 112(4), 1083–1088. 10.1073/pnas.1414894112

Fifer, J. E., Yasuda, N., Yamakita, T., Bove, C. B., & Davies, S. W. (2022). Genetic divergence and range expansion in a western North Pacific coral. Science of The Total Environment, 813, 152423. 10.1016/j.scitotenv.2021.152423

Frankham, R. (1995). CONSERVATION GENETICS. In AnnlL Rev. Genetics (Vol. 29). www.annualreviews.org

Franklin, I. R. (1980). Evolutionary Change in Small Populations. In n Conservation Biology: An Evolutionary-Ecological Perspective (pp. 35–49).

Freedman, D. A. (2007). Statistical models for causation. The SAGE Handbook of Social Science Methodology, 127–146.

Fuller, Z. L., Mocellin, V. J. L., Morris, L. A., Cantin, N., Shepherd, J., Sarre, L., Peng, J., Liao, Y., Pickrell, J., Andolfatto, P., Matz, M., Bay, L. K., & Przeworski, M. (2020). Population genetics of the coroal Acropora millepora: Toward genomic prediction of bleaching. Science, 369(6501). 10.1126/science.aba4674

Gignoux-Wolfsohn, S. A., Pinsky, M. L., Kerwin, K., Herzog, C., Hall, M., Bennett, A. B., Fefferman, N. H., & Maslo, B. (2021). Genomic signatures of selection in bats surviving white-nose syndrome. Molecular Ecology, 30(22), 5643–5657. 10.1111/mec.15813

Gilmore, T. D., & Wolenski, F. S. (2012). NF-κB: Where did it come from and why? In Immunological Reviews (Vol. 246, Issue 1, pp. 14–35). 10.1111/j.1600-065X.2012.01096.x

Gurgel, C. F. D., Camacho, O., Minne, A. J. P., Wernberg, T., & Coleman, M. A. (2020). Marine Heatwave Drives Cryptic Loss of Genetic Diversity in Underwater Forests. Current Biology, 30(7), 1199–1206.e2. 10.1016/j.cub.2020.01.051

Hansen, M. M., Olivieri, I., Waller, D. M., & Nielsen, E. E. (2012). Monitoring adaptive genetic responses to environmental change. Molecular Ecology, 21(6), 1311–1329. 10.1111/j.1365-294X.2011.05463.x

Harmelin-Vivien, M. L., & Laboute, P. (1986). Catastrophic impact of hurricanes on atoll outer reef slopes in the Tuamotu (French Polynesia). In Coral Reefs (Vol. 5).

Hedgecock, D. (1994). Temporal and spatial genetic structure of marine animal populations in the California Current. California Cooperative Oceanic Fisheries Investigations Reports, 35, 73–81.

Hoegh-Guldberg, O., Mumby, P. J., Hooten, A. J., Steneck, R. S., Greenfield, P., Gomez, E., Harvell, C. D., Sale, P. F., Edwards, A. J., Caldeira, K., Knowlton, N., Eakin, C. M., Iglesias-Prieto, R., Muthiga, N., Bradbury, R. H., Dubi, A., & Hatziolos, M. E. (2007). Coral Reefs Under Rapid Climate Change and Ocean Acidification. Science, 318(5857), 1737–1742. 10.1126/science.1152509

Holbrook, S. J., Adam, T. C., Edmunds, P. J., Schmitt, R. J., Carpenter, R. C., Brooks, A. J., Lenihan, H. S., & Briggs, C. J. (2018). Recruitment Drives Spatial Variation in Recovery Rates of Resilient Coral Reefs. Scientific Reports, 8(1). 10.1038/s41598-018-25414-8

Holland, O. J., Toomey, M., Ahrens, C., Hoffmann, A. A., Croft, L. J., Sherman, C. D. H., & Miller, A. D. (2022). Whole genome resequencing reveals signatures of rapid selection in a virus-affected commercial fishery. Molecular Ecology, 31(13), 3658–3671. 10.1111/mec.16499

Hormozdiari, F., Kostem, E., Kang, E. Y., Pasaniuc, B., & Eskin, E. (2014). Identifying causal variants at loci with multiple signals of association. ACM BCB 2014 - 5th ACM Conference on Bioinformatics, Computational Biology, and Health Informatics, 610–611. 10.1145/2649387.2660800

Howe-Kerr, L. I., Bachelot, B., Wright, R. M., Kenkel, C. D., Bay, L. K., & Correa, A. M. S. (2020). Symbiont community diversity is more variable in corals that respond poorly to stress. Global Change Biology, 26(4), 2220–2234. 10.1111/gcb.14999

Howells, E. J., Bauman, A. G., Vaughan, G. O., Hume, B. C. C., Voolstra, C. R., & Burt, J. A. (2020). Corals in the hottest reefs in the world exhibit symbiont fidelity not flexibility. Molecular Ecology, 29(5), 899–911. 10.1111/mec.15372

Howells, E. J., Bay, L. K., & Bay, R. A. (2022). Identifying, Monitoring, and Managing Adaptive Genetic Variation in Reef-Building Corals under Rapid Climate Warming (pp. 55– 70). 10.1007/978-3-031-07055-6_4

Hume, B. C. C., Mejia-Restrepo, A., Voolstra, C. R., & Berumen, M. L. (2020). Fine-scale delineation of Symbiodiniaceae genotypes on a previously bleached central Red Sea reef system demonstrates a prevalence of coral host-specific associations. Coral Reefs, 39(3), 583–601. 10.1007/s00338-020-01917-7

Hume, B. C. C., Smith, E. G., Ziegler, M., Warrington, H. J. M., Burt, J. A., LaJeunesse, T. C., Wiedenmann, J., & Voolstra, C. R. (2019). SymPortal: A novel analytical framework and platform for coral algal symbiont next-generation sequencing ITS2 profiling. Molecular Ecology Resources, 19(4), 1063–1080. 10.1111/1755-0998.13004

Hume, B. C. C., Ziegler, M., Poulain, J., Pochon, X., Romac, S., Boissin, E., de Vargas, C., Planes, S., Wincker, P., & Voolstra, C. R. (2018). An improved primer set and amplification protocol with increased specificity and sensitivity targeting the Symbiodinium ITS2 region. PeerJ, 2018(5). 10.7717/peerj.4816

Jones, A., & Berkelmans, R. (2010). Potential costs of acclimatization to a warmer climate: Growth of a reef coral with heat tolerant vs. sensitive symbiont types. PLoS ONE, 5(5). 10.1371/journal.pone.0010437

Jones, A. M., Berkelmans, R., Van Oppen, M. J. H., Mieog, J. C., & Sinclair, W. (2008). A community change in the algal endosymbionts of a scleractinian coral following a natural bleaching event: Field evidence of acclimatization. Proceedings of the Royal Society B: Biological Sciences, 275(1641), 1359–1365. 10.1098/rspb.2008.0069

Jun, G., Wing, M. K., Abecasis, G. R., & Kang, H. M. (2015). An efficient and scalable analysis framework for variant extraction and refinement from population-scale DNA sequence data. Genome Research, 25(6), 918–925. 10.1101/gr.176552.114

Käsbauer, T., Towb, P., Alexandrova, O., David, C. N., Dall’Armi, E., Staudigl, A., Stiening, B., & Böttger, A. (2007). The Notch signaling pathway in the cnidarian Hydra. Developmental Biology, 303(1), 376–390. 10.1016/j.ydbio.2006.11.022

Kayal, M., Vercelloni, J., Lison de Loma, T., Bosserelle, P., Chancerelle, Y., Geoffroy, S., Stievenart, C., Michonneau, F., Penin, L., Planes, S., & Adjeroud, M. (2012). Predator Crown-of-Thorns Starfish (Acanthaster planci) Outbreak, Mass Mortality of Corals, and Cascading Effects on Reef Fish and Benthic Communities. PLoS ONE, 7(10). 10.1371/journal.pone.0047363

Kemp, D. W., Hoadley, K. D., Lewis, A. M., Wham, D. C., Smith, R. T., Warner, M. E., & Lajeunesse, T. C. (2023). Thermotolerant coral-algal mutualisms maintain high rates of nutrient transfer while exposed to heat stress. Proceedings of the Royal Society B: Biological Sciences, 290(2007). 10.1098/rspb.2023.1403

Kirk, N. L., Howells, E. J., Abrego, D., Burt, J. A., & Meyer, E. (2018). Genomic and transcriptomic signals of thermal tolerance in heat-tolerant corals (Platygyra daedalea) of the Arabian/Persian Gulf. Molecular Ecology, 27(24), 5180–5194. 10.1111/mec.14934

Klingbeil, W. H., Montecinos, G. J., & Alberto, F. (2022). Giant kelp genetic monitoring before and after disturbance reveals stable genetic diversity in Southern California. Frontiers in Marine Science, 9. 10.3389/fmars.2022.947393

Klueter, A., Trapani, J., Archer, F. I., McIlroy, S. E., & Coffroth, M. A. (2017). Comparative growth rates of cultured marine dinoflagellates in the genus Symbiodinium and the effects of temperature and light. PLoS ONE, 12(11). 10.1371/journal.pone.0187707

Knowlton, N., Brainard, R. E., Fisher, R., Moews, M., Plaisance, L., & Caley, M. J. (2010). Chapter 4 Coral Reef Biodiversity.

Korneliussen, T. S., Albrechtsen, A., & Nielsen, R. (2014). ANGSD: Analysis of Next Generation Sequencing Data. BMC Bioinformatics, 15(1), 1–13. 10.1186/s12859-014-0356-4

Kriefall, N. G., Kanke, M. R., Aglyamova, G. V., & Davies, S. W. (2022). Reef environments shape microbial partners in a highly connected coral population. Proceedings of the Royal Society B: Biological Sciences, 289(1967). 10.1098/rspb.2021.2459

LaJeunesse, T. C., Parkinson, J. E., Gabrielson, P. W., Jeong, H. J., Reimer, J. D., Voolstra, C. R., & Santos, S. R. (2018). Systematic Revision of Symbiodiniaceae Highlights the Antiquity and Diversity of Coral Endosymbionts. Current Biology, 28(16), 2570–2580.e6. 10.1016/j.cub.2018.07.008

Langmead, B., & Salzberg L., S. (2013). Fast gapped-read alignment with Bowtie 2. Nature Methods, 9(4), 357–359. 10.1038/nmeth.1923.Fast

Leichter, J. J., Alldredge, A. L., Bernardi, G., Brooks, A. J., Carlson, C. A., Carpenter, R. C., Edmunds, P. J., Fewings, M. R., Hanson, K. M., Hench, J. L., Holbrook, S. J., Nelson, C. E., Schmitt, R. J., Toonen, R. J., Washburn, L., & Wyatt, A. S. J. (2013). Biological and physical interactions on a tropical island coral reef: Transport and retention processes on moorea, French Polynesia. Oceanography, 26(3), 52–63. 10.5670/oceanog.2013.45

Leichter, J. J., Seydel, K., & Gotschalk C. (2019). MCR LTER: Coral Reef: Benthic Water Temperature, ongoing since 2005 ver 12. In Environmental Data Initiative.

Leinbach, S. E., Speare, K. E., & Strader, M. E. (2023). Reef habitats structure symbiotic microalgal assemblages in corals and contribute to differential heat stress responses. Coral Reefs, 42(1), 205–217. 10.1007/s00338-022-02316-w

Lewis, R. E., Davy, S. K., Gardner, S. G., Rongo, T., Suggett, D. J., & Nitschke, M. R. (2022). Colony self-shading facilitates Symbiodiniaceae cohabitation in a South Pacific coral community. Coral Reefs, 41(5), 1433–1447. 10.1007/s00338-022-02292-1

Li, H., Handsaker, B., Wysoker, A., Fennell, T., Ruan, J., Homer, N., Marth, G., Abecasis, G., & Durbin, R. (2009). The Sequence Alignment/Map format and SAMtools. Bioinformatics, 25(16), 2078–2079. 10.1093/bioinformatics/btp352

Liu, H., Stephens, T. G., González-Pech, R. A., Beltran, V. H., Lapeyre, B., Bongaerts, P., Cooke, I., Aranda, M., Bourne, D. G., Forêt, S., Miller, D. J., van Oppen, M. J. H., Voolstra, C. R., Ragan, M. A., & Chan, C. X. (2018). Symbiodinium genomes reveal adaptive evolution of functions related to coral-dinoflagellate symbiosis. Communications Biology, 1, 95. 10.1038/s42003-018-0098-3

Liu, X., & Fu, Y. X. (2020). Stairway Plot 2: demographic history inference with folded SNP frequency spectra. Genome Biology, 21(1), 1–9. 10.1186/s13059-020-02196-9

López-Nandam, E. H., Albright, R., Hanson, E. A., Sheets, E. A., & Palumbi, S. R. (2023). Mutations in coral soma and sperm imply lifelong stem cell renewal and cell lineage selection. Proceedings of the Royal Society B: Biological Sciences, 290(1991). 10.1098/rspb.2022.1766

Loya, Y., Sakai, K., Yamazato, K., Nakano, Y., Sambali, H., van Woesik, R., Loya, Sakai, Yamazato, Nakano, Sambali, & Van, W. (2001). Coral bleaching: the winners and the losers. Ecology Letters, 4(2), 122–131. 10.1046/j.1461-0248.2001.00203.x

Marchese, A., Sawzdargo, M., Nguyen, T., Cheng, R., Heng, H. H. Q., Nowak, T., Im, D.-S., Lynch, K. R., George, S. R., & O’dowd, B. F. (1999). Discovery of Three Novel Orphan G-Protein-Coupled Receptors. http://www.idealibrary.com

Marlow, H., Roettinger, E., Boekhout, M., & Martindale, M. Q. (2012). Functional roles of Notch signaling in the cnidarian Nematostella vectensis. Developmental Biology, 362(2), 295–308. 10.1016/j.ydbio.2011.11.012

Mason, B., Koyanagi, M., Sugihara, T., Iwasaki, M., Slepak, V., Miller, D. J., Sakai, Y., & Terakita, A. (2023). Multiple opsins in a reef-building coral, Acropora millepora. Scientific Reports, 13(1). 10.1038/s41598-023-28476-5

Mason, B., Schmale, M., Gibbs, P., Miller, M. W., Wang, Q., Levay, K., Shestopalov, V., & Slepak, V. Z. (2012). Evidence for Multiple Phototransduction Pathways in a Reef-Building Coral. PLoS ONE, 7(12). 10.1371/journal.pone.0050371

Matz, M. V., Treml, E. A., Aglyamova, G. V., & Bay, L. K. (2018). Potential and limits for rapid genetic adaptation to warming in a Great Barrier Reef coral. PLoS Genetics, 14(4), 1–27. 10.1371/journal.pgen.1007220

Matz, M. V., Treml, E. A., & Haller, B. C. (2020). Estimating the potential for coral adaptation to global warming across the Indo-West Pacific. Global Change Biology, 26(6), 3473–3481. 10.1111/gcb.15060

Morikawa, M. K., & Palumbi, S. R. (2019). Using naturally occurring climate resilient corals to construct bleaching-resistant nurseries. Proceedings of the National Academy of Sciences of the United States of America, 116(21), 10586–10591. 10.1073/pnas.1721415116

Münder, S., Käsbauer, T., Prexl, A., Aufschnaiter, R., Zhang, X., Towb, P., & Böttger, A. (2010). Notch signalling defines critical boundary during budding in Hydra. Developmental Biology, 344(1), 331–345. 10.1016/j.ydbio.2010.05.517

Nei, M., Maruyama, T., & Chakraborty, R. (1975). The Bottleneck Effect and Genetic Variability in Populations (Vol. 29, Issue 1). https://www.jstor.org/stable/2407137

Okansen, J., Blanchet, F. G., Friendly, M., Kindt, R., Legendre, P., McGlinn, D., Minchin, P. R., O’Hara, R. B., Simpson, G. L., & Solymos, P. (2020). vegan: community ecology package. R package version 2.5–7.

Oliver, T. A., & Palumbi, S. R. (2011). Many corals host thermally resistant symbionts in high-temperature habitat. Coral Reefs, 30(1), 241–250. 10.1007/s00338-010-0696-0

Penin, L., Adjeroud, M., Schrimm, M., & Lenihan, H. S. (2007). High spatial variability in coral bleaching around Moorea (French Polynesia): patterns across locations and water depths. Comptes Rendus - Biologies, 330(2), 171–181. 10.1016/j.crvi.2006.12.003 Picard Toolkit. (2019). http://broadinstitute.github.io/picard.

Pilczynska, J., Cocito, S., Boavida, J., Serrão, E., & Queiroga, H. (2016). Genetic diversity and local connectivity in the Mediterranean red gorgonian coral after mass mortality events. PLoS ONE, 11(3). 10.1371/journal.pone.0150590

Pratchett, M. S., McCowan, D., Maynard, J. A., & Heron, S. F. (2013). Changes in Bleaching Susceptibility among Corals Subject to Ocean Warming and Recurrent Bleaching in Moorea, French Polynesia. PLoS ONE, 8(7). 10.1371/journal.pone.0070443

Pujolar, J. M., Vincenzi, S., Zane, L., Jesensek, D., de Leo, G. A., & Crivelli, A. J. (2011). The effect of recurrent floods on genetic composition of marble trout populations. PLoS ONE, 6(9). 10.1371/journal.pone.0023822

Quigley, K. M., & van Oppen, M. J. H. (2022). Predictive models for the selection of thermally tolerant corals based on offspring survival. Nature Communications, 13(1). 10.1038/s41467-022-28956-8

R Core Team. (2023). R: A language and environment for statistical computing. R Foundation for Statistical Computing.

Radwan, J., Biedrzycka, A., & Babik, W. (2010). Does reduced MHC diversity decrease viability of vertebrate populations? In Biological Conservation (Vol. 143, Issue 3, pp. 537–544). 10.1016/j.biocon.2009.07.026

Rasmussen, M. S., Garcia-Erill, G., Korneliussen, T. S., Wiuf, C., & Albrechtsen, A. (2022). Estimation of site frequency spectra from low-coverage sequencing data using stochastic EM reduces overfitting, runtime, and memory usage. Genetics, 222(4). 10.1093/genetics/iyac148

Reid, B. N., Star, B., & Pinsky, M. L. (2023). Detecting parallel polygenic adaptation to novel evolutionary pressure in wild populations: A case study in Atlantic cod (Gadus morhua). Philosophical Transactions of the Royal Society B: Biological Sciences, 378(1881). 10.1098/rstb.2022.0190

Ripley, B., Venables, B., Bates, D. M., Hornik, K., Gebhardt, A., Firth, D., & Ripley, M. B. (2013). Package ‘mass.’ Cran r, 538, 113–120.

Rose, N. H., Bay, R. A., Morikawa, M. K., & Palumbi, S. R. (2018). Polygenic evolution drives species divergence and climate adaptation in corals. Evolution, 72(1), 82–94. 10.1111/evo.13385

Rose, N. H., Bay, R. A., Morikawa, M. K., Thomas, L., Sheets, E. A., & Palumbi, S. R. (2021). Genomic analysis of distinct bleaching tolerances among cryptic coral species. Proceedings of the Royal Society B: Biological Sciences, 288(1960). 10.1098/rspb.2021.0678

Rose, N. H., Seneca, F. O., & Palumbi, S. R. (2015). Gene Networks in the Wild : Identifying Transcriptional. Genome Biology and Evolution, 8(1), 243–252. 10.5061/dryad.and

Schiebelhut, L. M., Puritz, J. B., & Dawson, M. N. (2018). Decimation by sea star wasting disease and rapid genetic change in a keystone species, Pisaster ochraceus. PNAS, 115(27), 7069–7074. 10.6071/M3WW84

Schneider, C. A., Rasband, W. S., & Eliceiri, K. W. (2012). NIH Image to ImageJ: 25 years of image analysis. In Nature Methods (Vol. 9, Issue 7, pp. 671–675). 10.1038/nmeth.2089

Seneca, F. O., & Palumbi, S. R. (2015). The role of transcriptome resilience in resistance of corals to bleaching. Molecular Ecology, 24(7), 1467–1484. 10.1111/mec.13125

Shoguchi, E., Shinzato, C., Kawashima, T., Gyoja, F., Mungpakdee, S., Koyanagi, R., Takeuchi, T., Hisata, K., Tanaka, M., Fujiwara, M., Hamada, M., Seidi, A., Fujie, M., Usami, T., Goto, H., Yamasaki, S., Arakaki, N., Suzuki, Y., Sugano, S., … Satoh, N. (2013). Draft assembly of the symbiodinium minutum nuclear genome reveals dinoflagellate gene structure. Current Biology, 23(15), 1399–1408. 10.1016/j.cub.2013.05.062

Siebeck, U. E., Marshall, N. J., Klüter, A., & Hoegh-Guldberg, O. (2006). Monitoring coral bleaching using a colour reference card. Coral Reefs, 25(3), 453–460. 10.1007/s00338-006-0123-8

Silva-Lima, A. W., Walter, J. M., Garcia, G. D., Ramires, N., Ank, G., Meirelles, P. M., Nobrega, A. F., Siva-Neto, I. D., Moura, R. L., Salomon, P. S., Thompson, C. C., & Thompson, F. L. (2015). Multiple Symbiodinium Strains Are Hosted by the Brazilian Endemic Corals Mussismilia spp. Microbial Ecology, 70(2), 301–310. 10.1007/s00248-015-0573-z

Silverstein, R. N., Cunning, R., & Baker, A. C. (2017). Tenacious D: Symbiodinium in clade D remain in reef corals at both high and low temperature extremes despite impairment. Journal of Experimental Biology, 220(7), 1192–1196. 10.1242/jeb.148239

Smith, E. G., Hazzouri, K. M., Choi, J. Y., Delaney, P., Al-Kharafi, M., Howells, E. J., Aranda, M., & Burt, J. A. (2022). Signatures of selection underpinning rapid coral adaptation to the world’s warmest reefs. In Sci. Adv (Vol. 8). https://www.science.org

Speare, K. E., Adam, T. C., Winslow, E. M., Lenihan, H. S., & Burkepile, D. E. (2022). Size dependent mortality of corals during marine heatwave erodes recovery capacity of a coral reef. Global Change Biology, 28(4), 1342–1358. 10.1111/gcb.16000

Tago, K., Funakoshi-Tago, M., Sakinawa, M., Mizuno, N., & Itoh, H. (2010). κB-Ras is a nuclear-cytoplasmic small GTpase that inhibits NF-κB activation through the suppression of transcriptional activation of p65/RelA. Journal of Biological Chemistry, 285(40), 30622– 30633. 10.1074/jbc.M110.117028

Thomas, L., Şahin, D., Adam, A. S., Grimaldi, C. M., Ryan, N. M., Duffy, S. L., Underwood, J. N., Kennington, W. J., & Gilmour, J. P. (2024). Resilience to periodic disturbances and the long-term genetic stability in Acropora coral. Communications Biology, 7(1), 410. 10.1038/s42003-024-06100-0

Traylor-Knowles, N., Rose, N. H., & Palumbi, S. R. (2017). The cell specificity of gene expression in the response to heat stress in corals. The Journal of Experimental Biology, March, jeb.155275. 10.1242/jeb.155275

Varasteh, T., Salazar, V., Tschoeke, D., Francini-Filho, R. B., Swings, J., Garcia, G., Thompson, C. C., & Thompson, F. L. (2022). Breviolum and Cladocopium Are Dominant Among Symbiodiniaceae of the Coral Holobiont Madracis decactis. Microbial Ecology, 84(2), 325–335. 10.1007/s00248-021-01868-8

Watterson, G. A. (1975). On the Number of Segregating Sites in Genetical Models without Recombination. In POPULATION BIOLOGY (Vol. 7).

Weis, V. M. (2019). Cell Biology of Coral Symbiosis: Foundational Study Can Inform Solutions to the Coral Reef Crisis. Integrative and Comparative Biology, 59(4), 845–855. 10.1093/icb/icz067

Zhang, J., Richards, Z. T., Adam, A. A. S., Chan, C. X., Shinzato, C., Gilmour, J., Thomas, L., Strugnell, J. M., Miller, D. J., & Cooke, I. (2022). Evolutionary Responses of a Reef-building Coral to Climate Change at the End of the Last Glacial Maximum. Molecular Biology and Evolution, 39(10). 10.1093/molbev/msac201

Zhou, X., Appl, A., & Author, S. (2017). A UNIFIED FRAMEWORK FOR VARIANCE COMPONENT ESTIMATION WITH SUMMARY STATISTICS IN GENOME-WIDE ASSOCIATION STUDIES 1 HHS Public Access Author manuscript. Ann Appl Stat, 11(4), 2027–2051. 10.1214/17-AOAS1052SUPP;.pdf

Zhou, X., & Stephens, M. (2012). Genome-wide efficient mixed-model analysis for association studies. Nature Genetics, 44(7), 821–824. 10.1038/ng.2310

Zhu, X., Stephens, M., Appl, A., Author, S., & Stat, A. A. (2018). BAYESIAN LARGE-SCALE MULTIPLE REGRESSION WITH SUMMARY STATISTICS FROM GENOME-WIDE ASSOCIATION STUDIES 1 HHS Public Access Author manuscript. 10.1214/17-AOAS1046SUPP;.pdf

