## Supplemental Material for "Genomic signatures of coral adaptation and recovery following a mass mortality event"

Supplementary Information

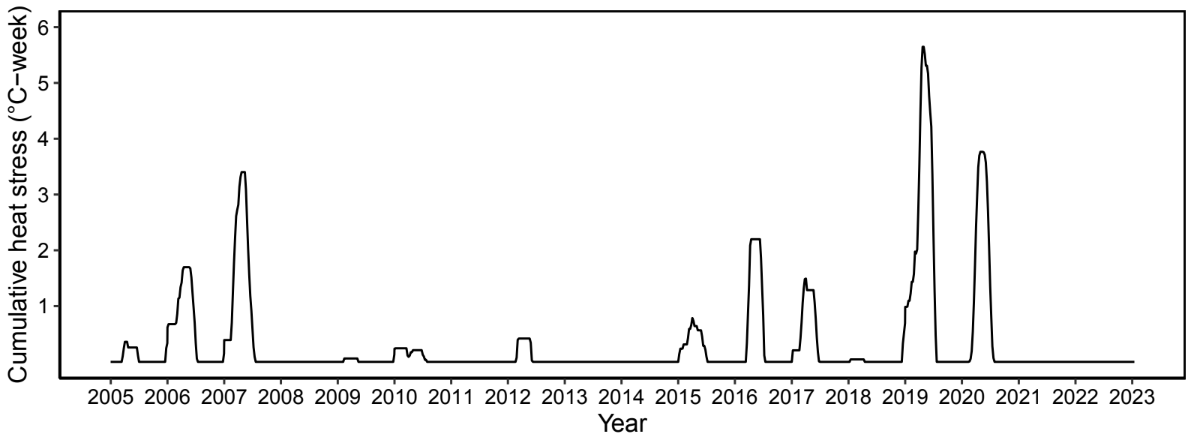
Figures

Supplementary Figure 1. Cumulative heat stress experienced across all MCR LTER sites (0-6) at 10 m depth on the forereef of Mo’orea over the prior 18 years. Cumulative heat stress was calculated as the 12-week running sum of all average weekly temperatures greater than 29°C. Data were collected as part of the MCR LTER core time series data collection (Edmunds & Moorea Coral Reef LTER, 2022).

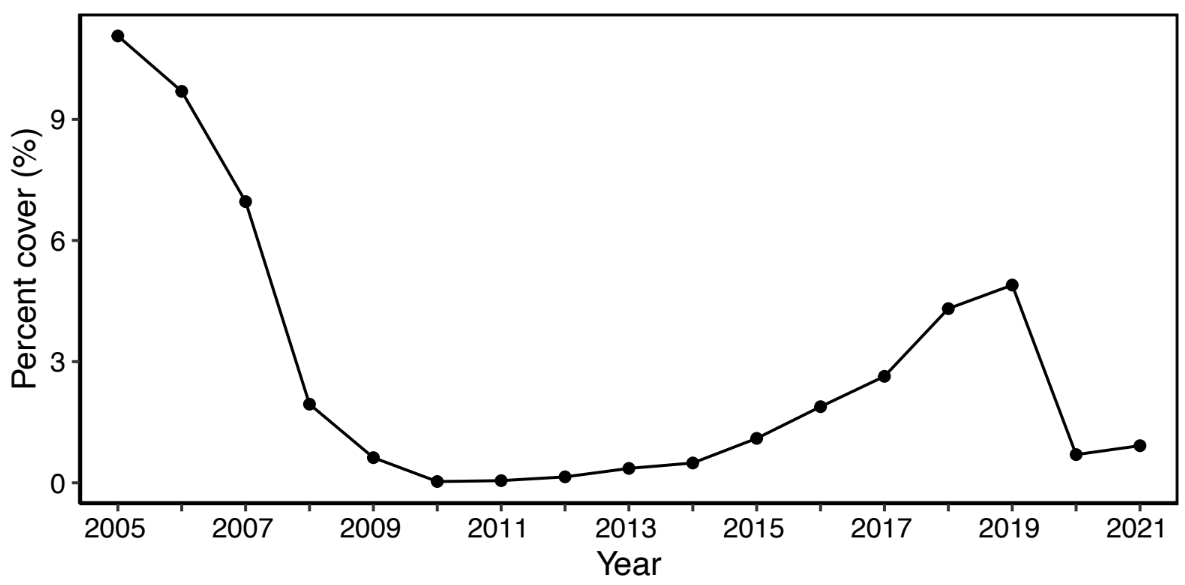
Supplementary Figure 2. *Acropora* spp. percent cover on the deep forereef (10 m depth) of Mo’orea, French Polynesia. Coverage surveys were conducted at MCR LTER sites 1-6 at 10 m water depth in April of each year. Each point represents the average across all sites per year. Data were collected as part of the MCR LTER core time series data collection (Edmunds & Moorea Coral Reef LTER, 2022).

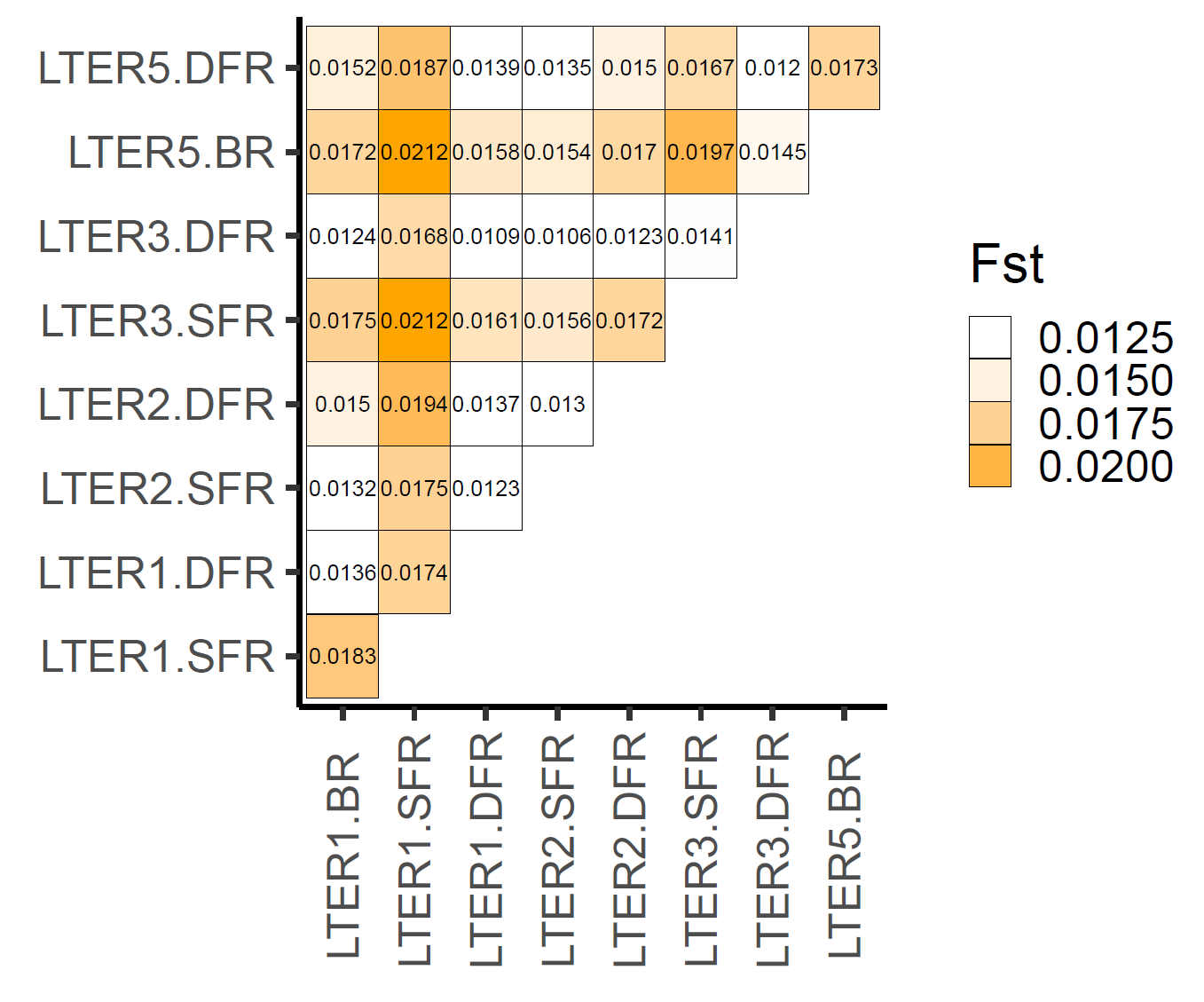
Supplemental Figure 3. Pre-mortality comparisons of pairwise *F*_ST_ between all sites, showing limited population structure. BR: Backreef; SFR: Shallow forereef; DFR: Deep forereef.

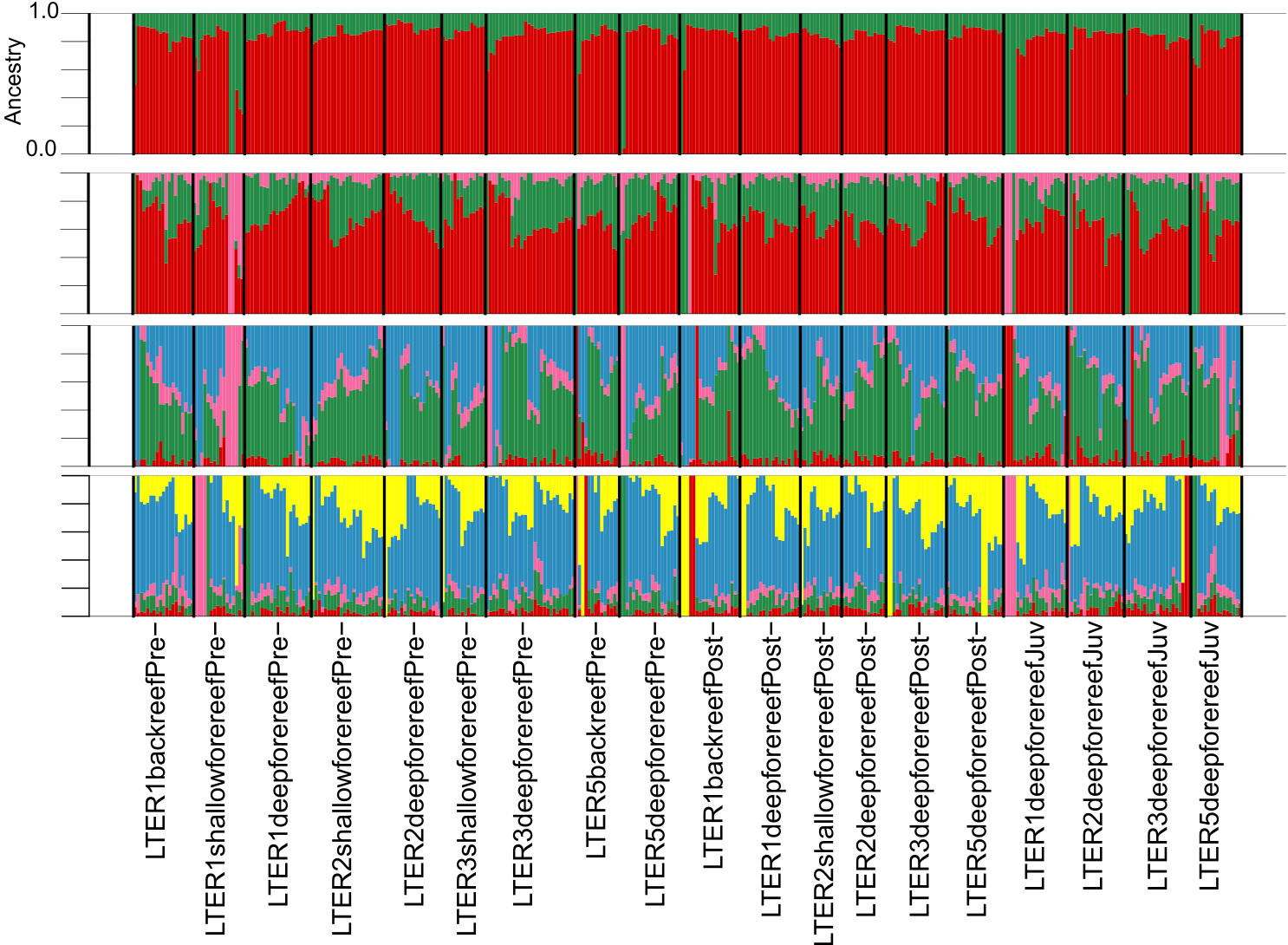
Supplemental Figure 4. Admixture plot showing all loci, Ks 2:5.

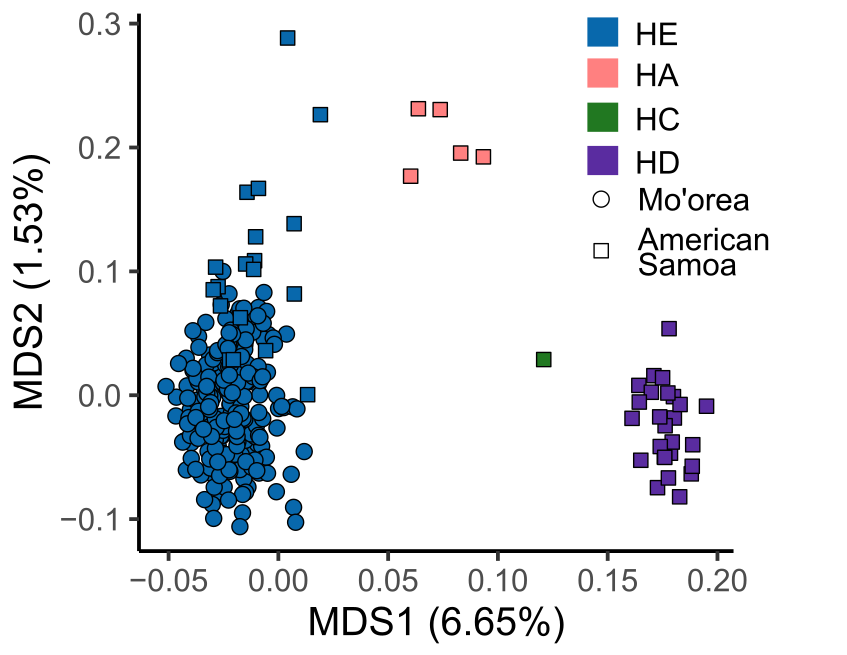

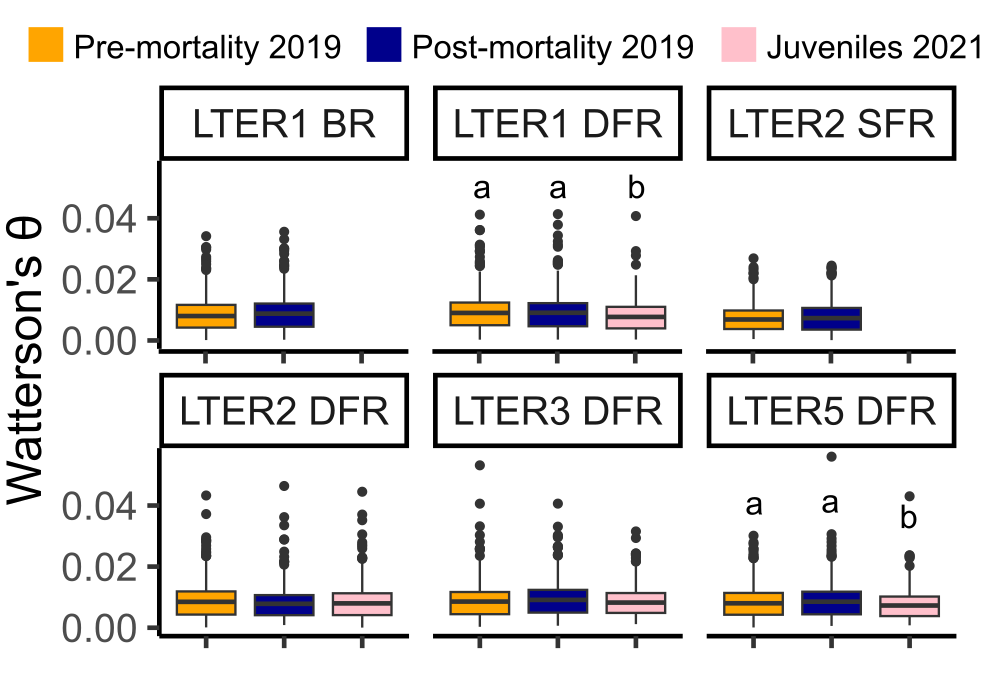
Supplemental Figure 5. MDS plot with *A. hyacinthus* WGS data from Rose et al. (2021), showing Mo’orea clustering with HE lineage on MDS1.

Supplemental Figure 6. Comparisons of nucleotide diversity (Watterson’s θ) between the overlapping sites for the three timepoints (pre- and post- mortality and juveniles). Letter denotes significance per Dunn’s test (1964) with a Benjamini-Hochberg multiple test correction. BR: Backreef; SFR: Shallow forereef; DFR: Deep forereef.

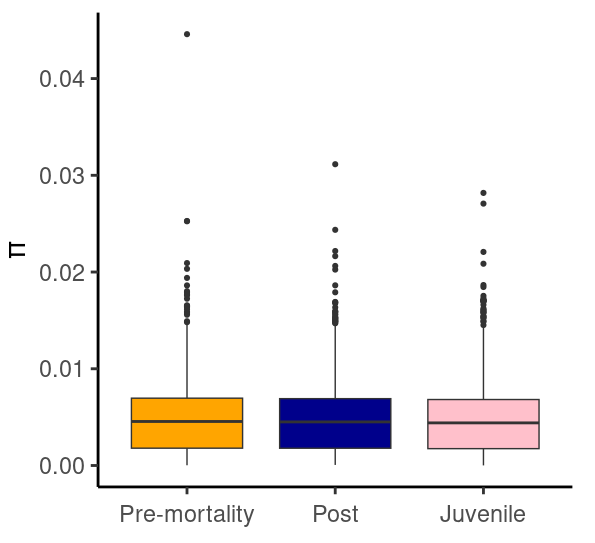

Supplemental Figure 7. π (pairwise theta in ANGSD) across scaffolds compared across the three timepoints (overlapping sites only). No letters denotes lack of significance per Dunn’s test (1964) with a Benjamini-Hochberg multiple test correction.

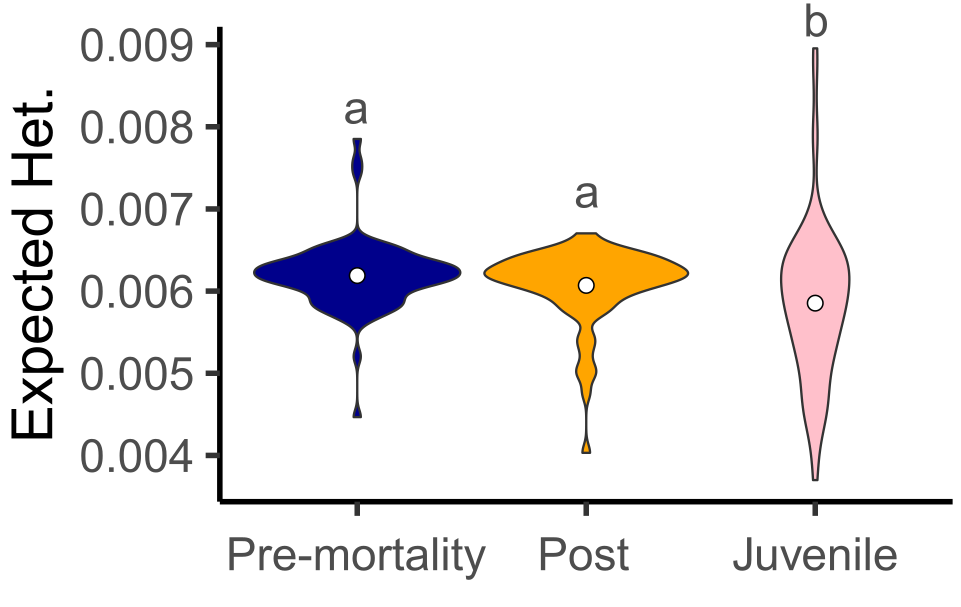

Supplemental Figure 8. Individual expected heterozygosity compared across the three timepoints (overlapping sites only). Letter denotes significance per Dunn’s test (1964) with a Benjamini-Hochberg multiple test correction.

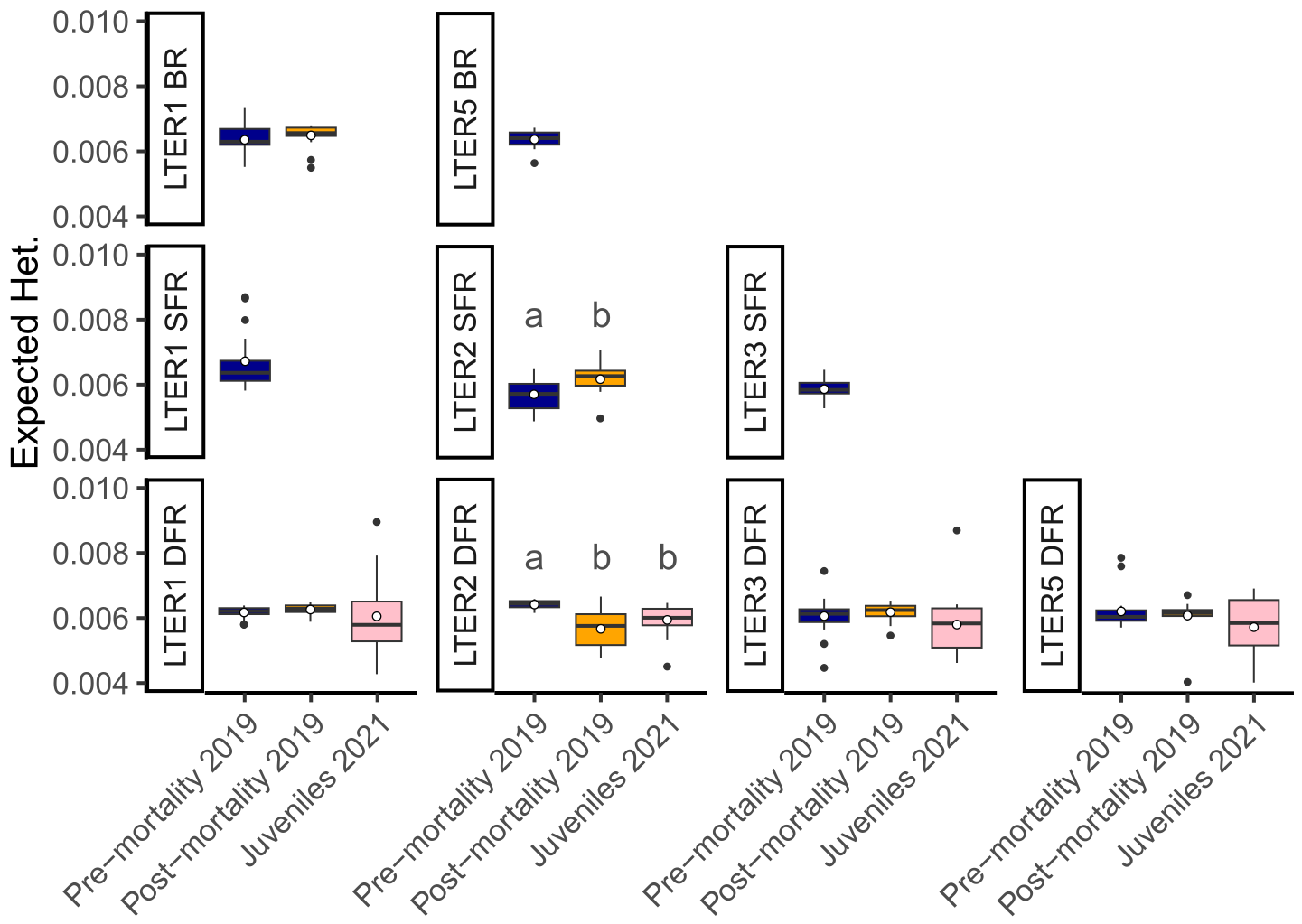

Supplemental Figure 9. Individual expected heterozygosity between all timepoints and site combinations sampled. Letters denote significance within each site per Dunn’s test (1964) with a Benjamini-Hochberg multiple test correction. Each facet that contains no letters had no significant differences.  BR: Backreef; SFR: Shallow forereef; DFR: Deep forereef.

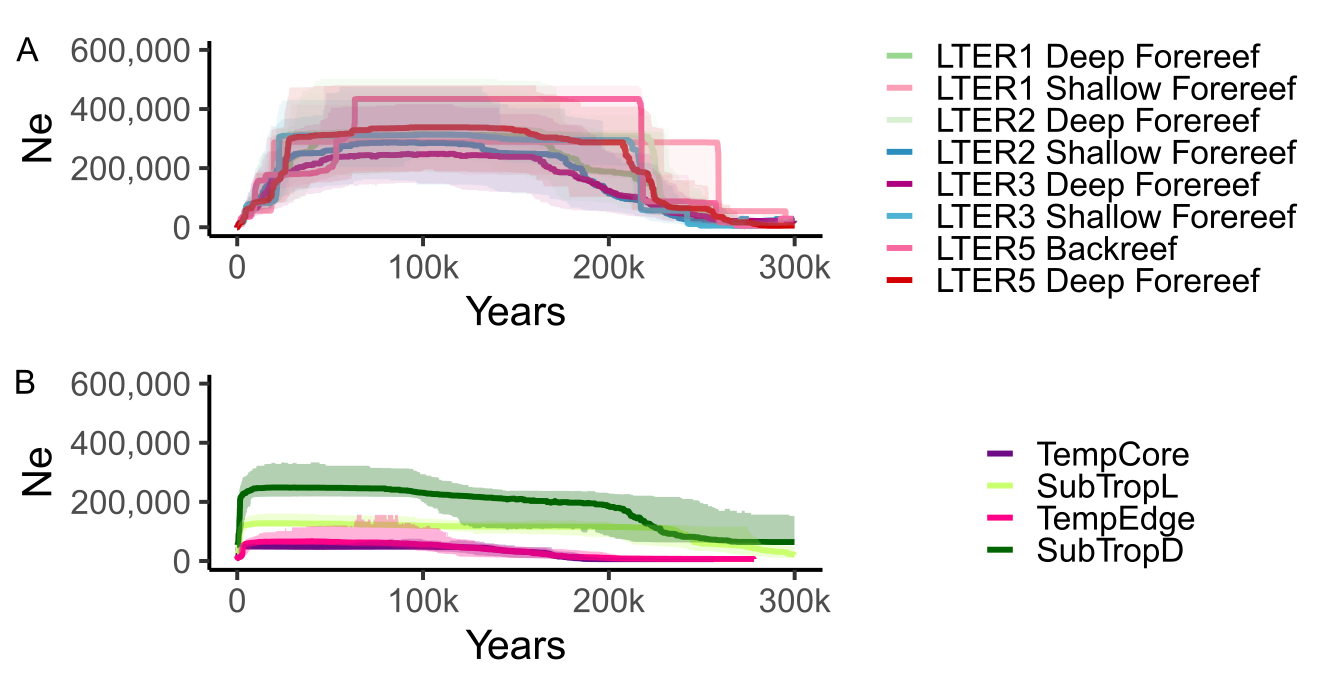

Supplemental Figure 10. Stairway plot showing Mo’orea *A. hyacinthus* has historically higher effective populations than Japan populations. A) Mo’orea populations from samples in May only. B) Japan populations representing three different lineages (SubTropL, SubTropD and Temp) of *A. hyacinthus* from Fifer et al. (2022)

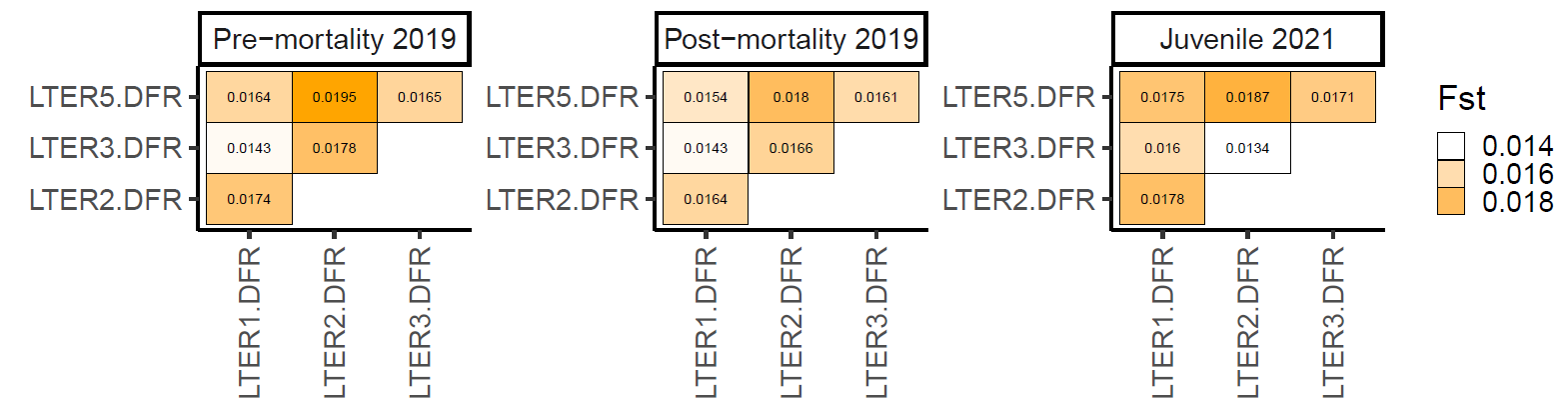

Supplemental Figure 11. Pairwise *F*_ST_ between deep forereef sites (DFR) for each timepoint.

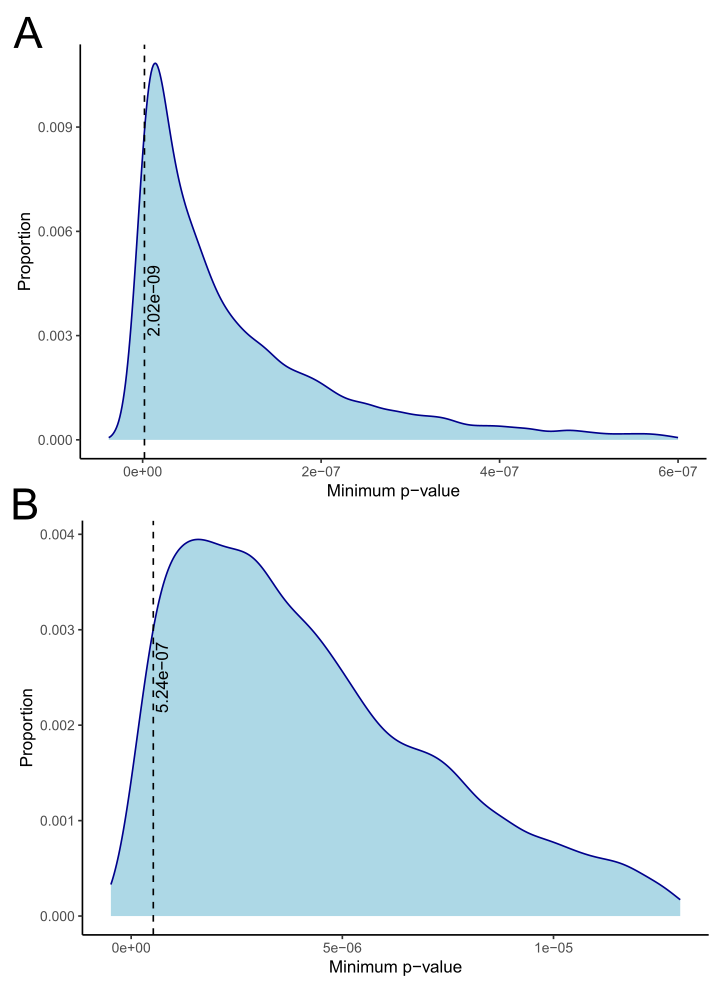

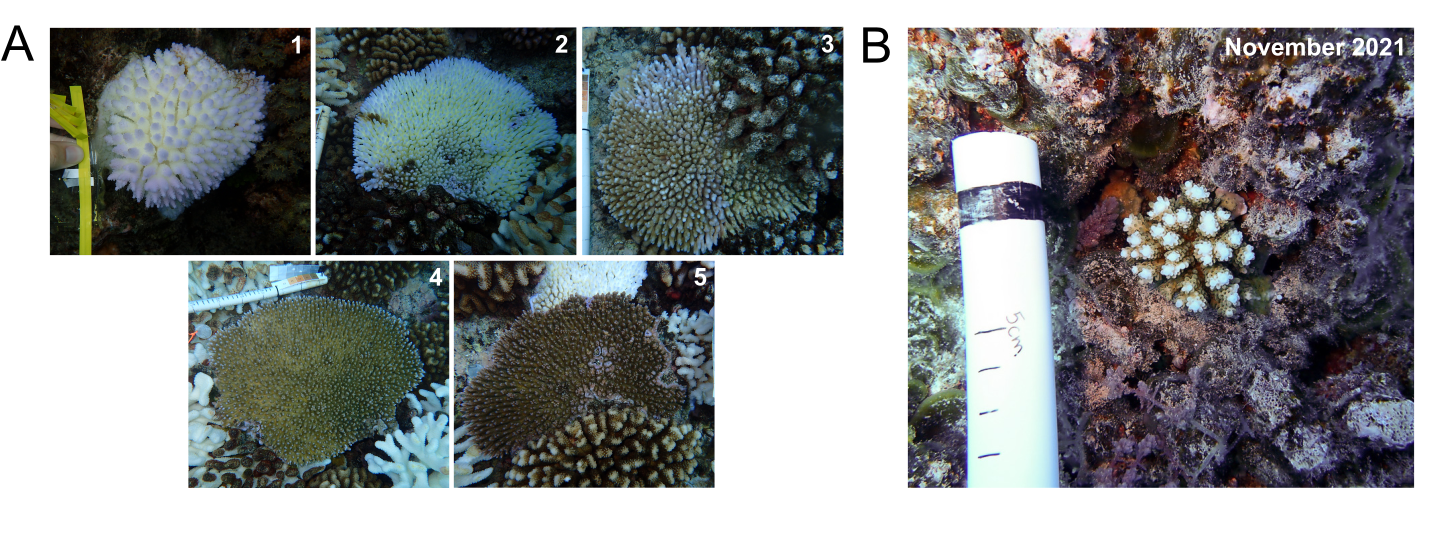

Supplemental Figure 12. Observed *A. hyacinthus* colonies during and after the 2019 thermal anomaly. A) Representative colonies illustrating the scale used to categorize bleaching intensity. Each colony was assigned an integer score from 1 to 5, with 1 indicating stark white bleaching and 5 indicating dark pigmentation and no bleaching. B) A juvenile *A. hyacinthus* colony settled after the 2019 MME, photographed in November 2021. Figure adapted from Leinbach et al. (2023).
Supplemental Figure 13. Distribution of minimal *p*-values determined from a permutation test to estimate a threshold of genome-wide significance for bleaching resilience (A) and bleaching survival (B). The 95^th^ percentile for each is indicated with a vertical line and was used as the genome-wide significance threshold.

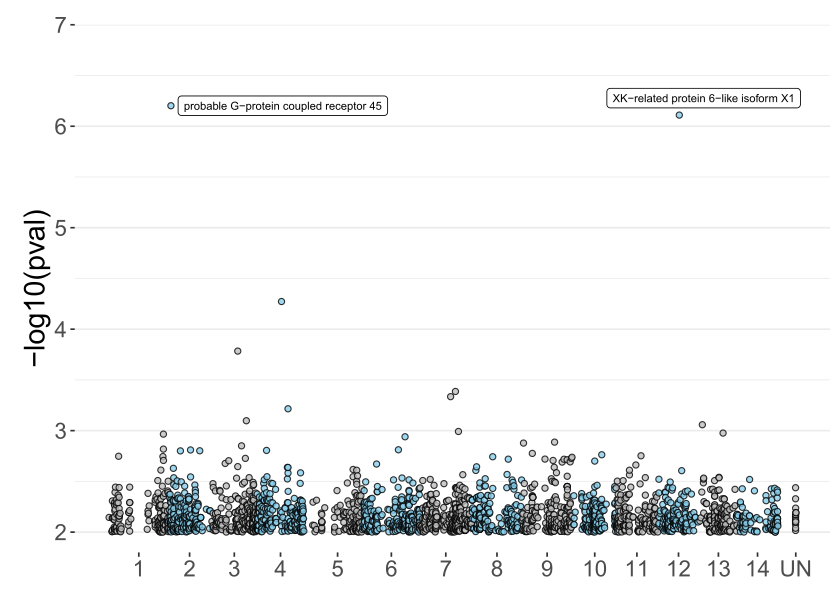

Supplemental Figure 14. Manhattan plot for bleaching survival GWAS performed on backreef samples only. Only loci with *p*-value < 0.01 are plotted here. Unplaced scaffolds were collapsed into one scaffold (UN).

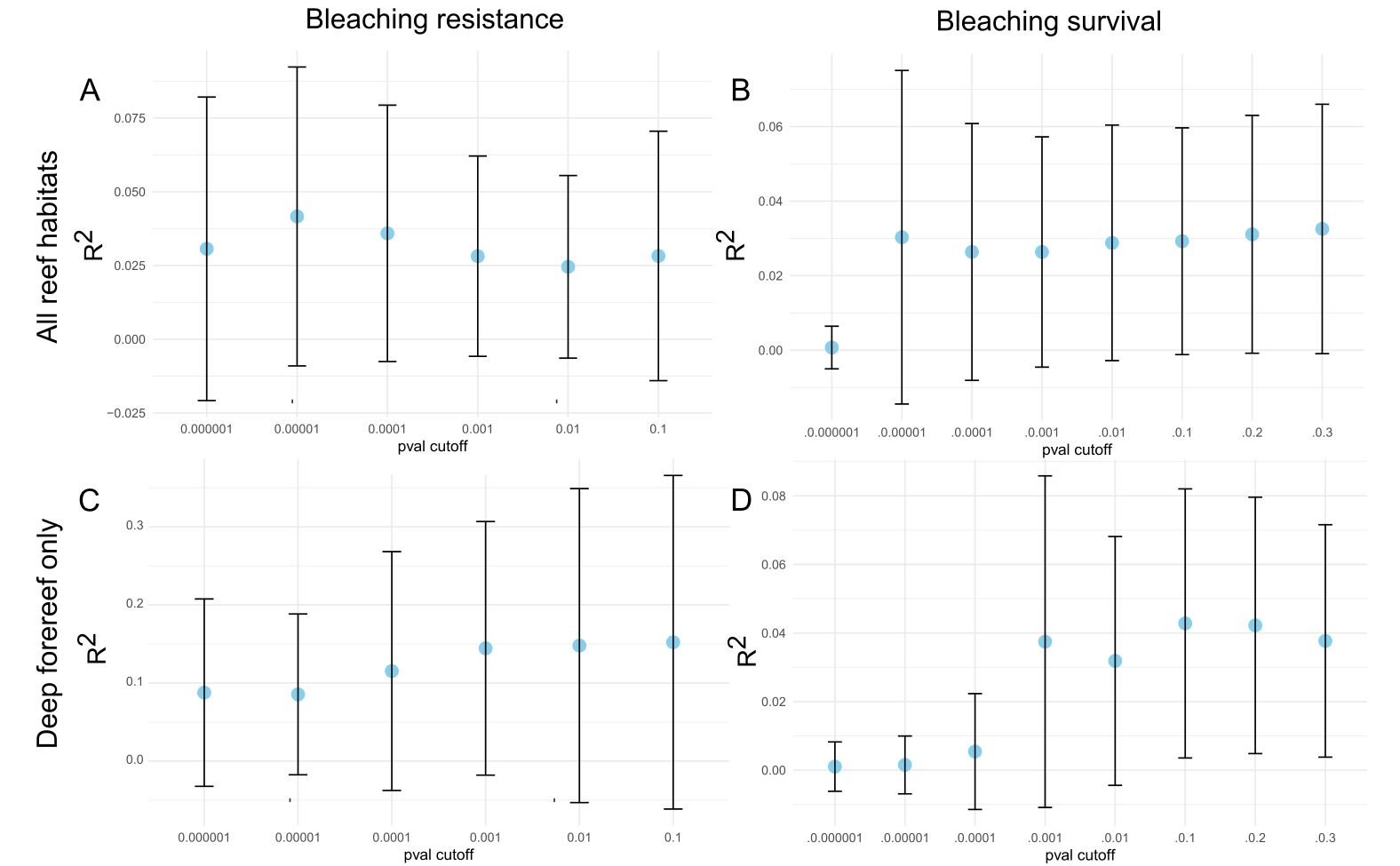

Supplemental Figure 15. Comparison of R^2^ for the 100 partitioned test sets at different GWAS *p*-value thresholds for PGS building. PGS built using backreef, shallow forereef and deep forereef habitats for A) pre-mortality samples taken during the bleaching event with health score as the trait and B) pre- and post-mortality samples with survival as the trait. PGS built using deep forereef habitat only for C) pre-mortality samples taken during the bleaching event with health score as the trait and D) pre- and post-mortality samples with survival as the trait. Plot shows mean and standard deviation of R^2^.

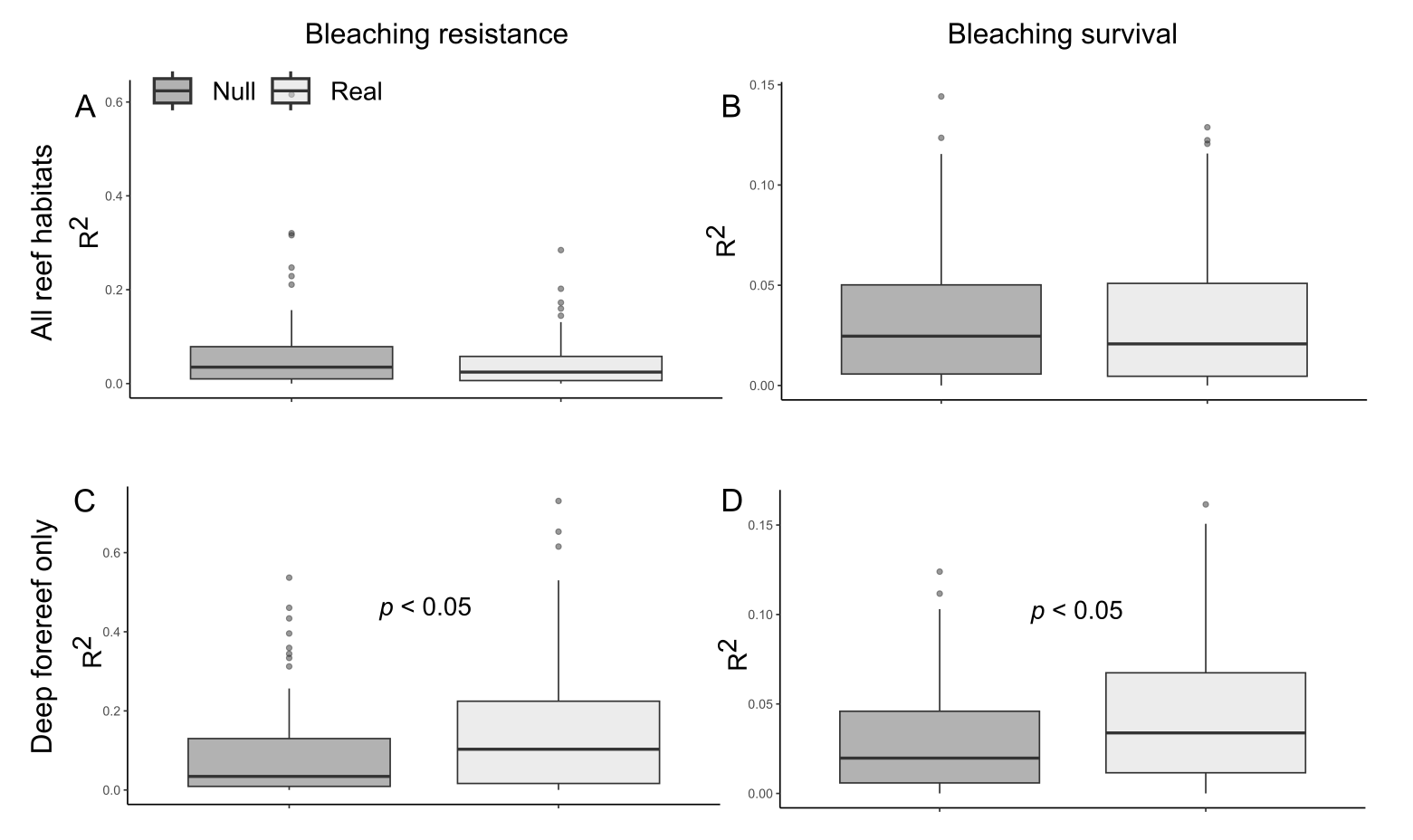

Supplemental Figure 16. Comparison of R^2^ for the 100 partitioned test sets between the real PGS (using optimal *p*-value threshold from Supp. Fig. 15) and a null PGS (randomly selected loci). PGS built using backreef, shallow forereef and deep forereef habitats for A) pre-mortality samples taken during the bleaching event with health score as the trait and B) pre- and post-mortality samples with survival as the trait. PGS built using deep forereef habitat only for C) pre-mortality samples taken during the bleaching event with health score as the trait and D) pre- and post-mortality samples with survival as the trait. Plot shows mean and standard deviation of R^2^.

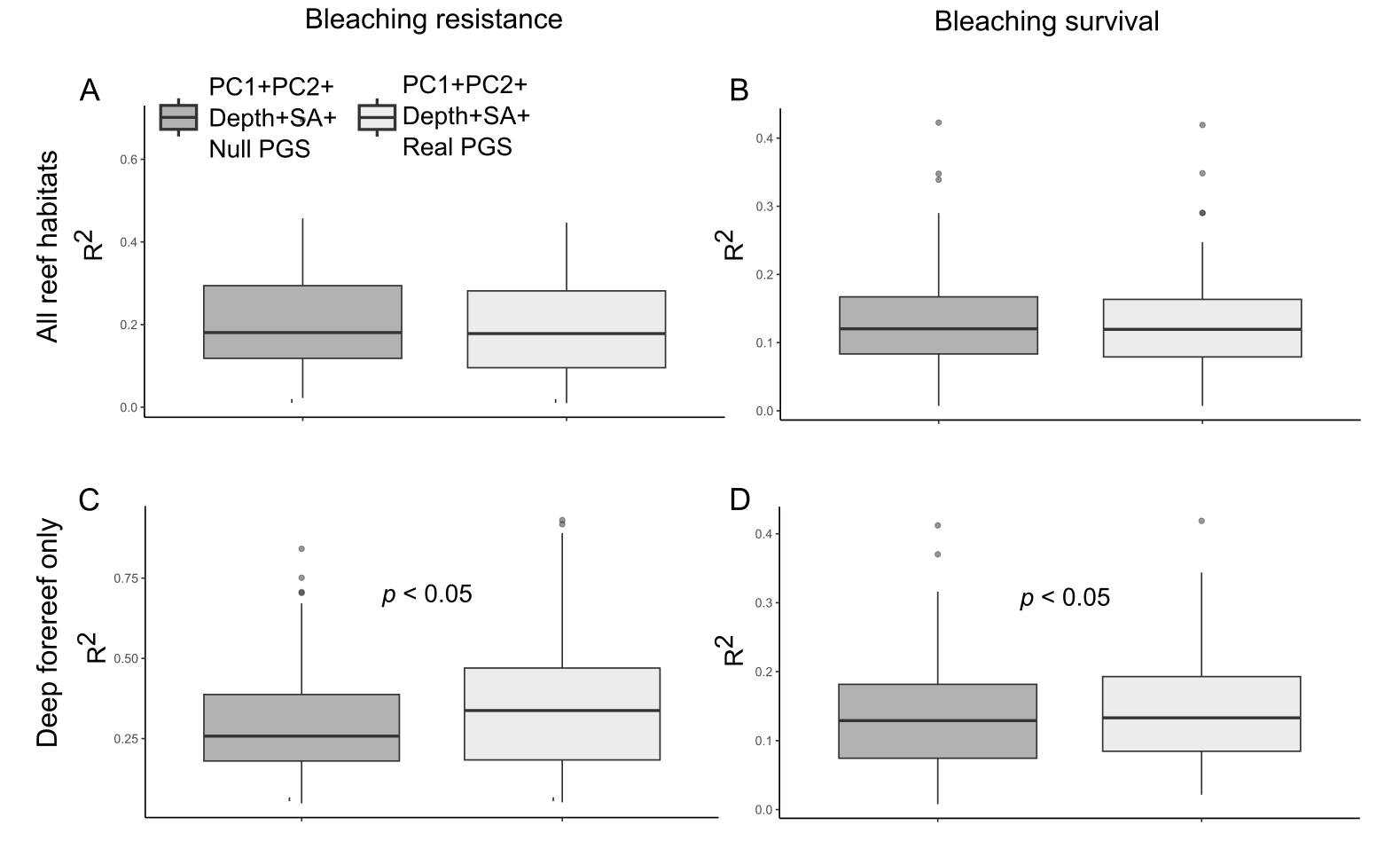

Supplemental Figure 17. Comparison of R^2^ for the 100 partitioned test sets between a model with all non-genetic covariates except symbiont proportions and the real PGS (using optimal *p*-value threshold from Supp. Fig. 15) versus a model with all non-genetic covariates excepting symbiont proportions and a null PGS (randomly selected loci). PGS built using backreef, shallow forereef and deep forereef habitats for A) pre-mortality samples taken during the bleaching event with health score as the trait and B) pre- and post-mortality samples with survival as the trait. PGS built using deep forereef habitat only for C) pre-mortality samples taken during the bleaching event with health score as the trait and D) pre- and post-mortality samples with survival as the trait. Plot shows mean and standard deviation of R^2^. SA: Surface area.

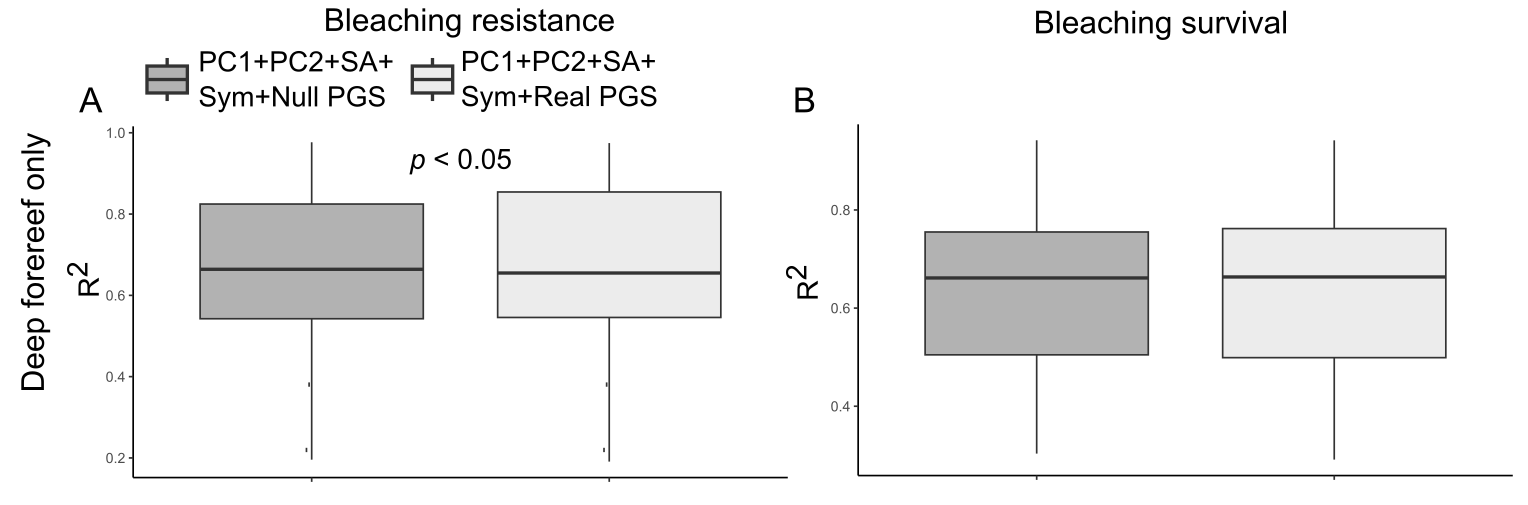

Supplemental Figure 18. Comparison of R^2^ for the 100 partitioned test sets between a model with all non-genetic covariates and the real PGS (using optimal *p*-value threshold from Supp. Fig.15) versus a model with all non-genetic covariates and a null PGS (randomly selected loci). PGS built using deep forereef habitat only for A) pre-mortality samples taken during the bleaching event with health score as the trait and B) pre- and post-mortality samples with survival as the trait. Plot shows mean and standard deviation of R^2^.

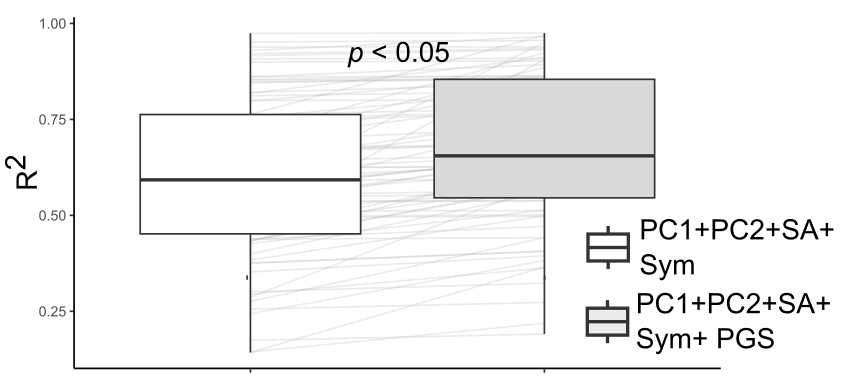

Supplemental Figure 19. Comparison of R^2^ for the 100 partitioned test sets between a model with all non-genetic covariates versus a model with all non-genetic covariates and the real PGS (using optimal *p*-value threshold from Supp. Fig. 15). PGS was built using deep forereef habitat only for pre-mortality samples taken during the bleaching event with health score as the trait. SA: Surface area.

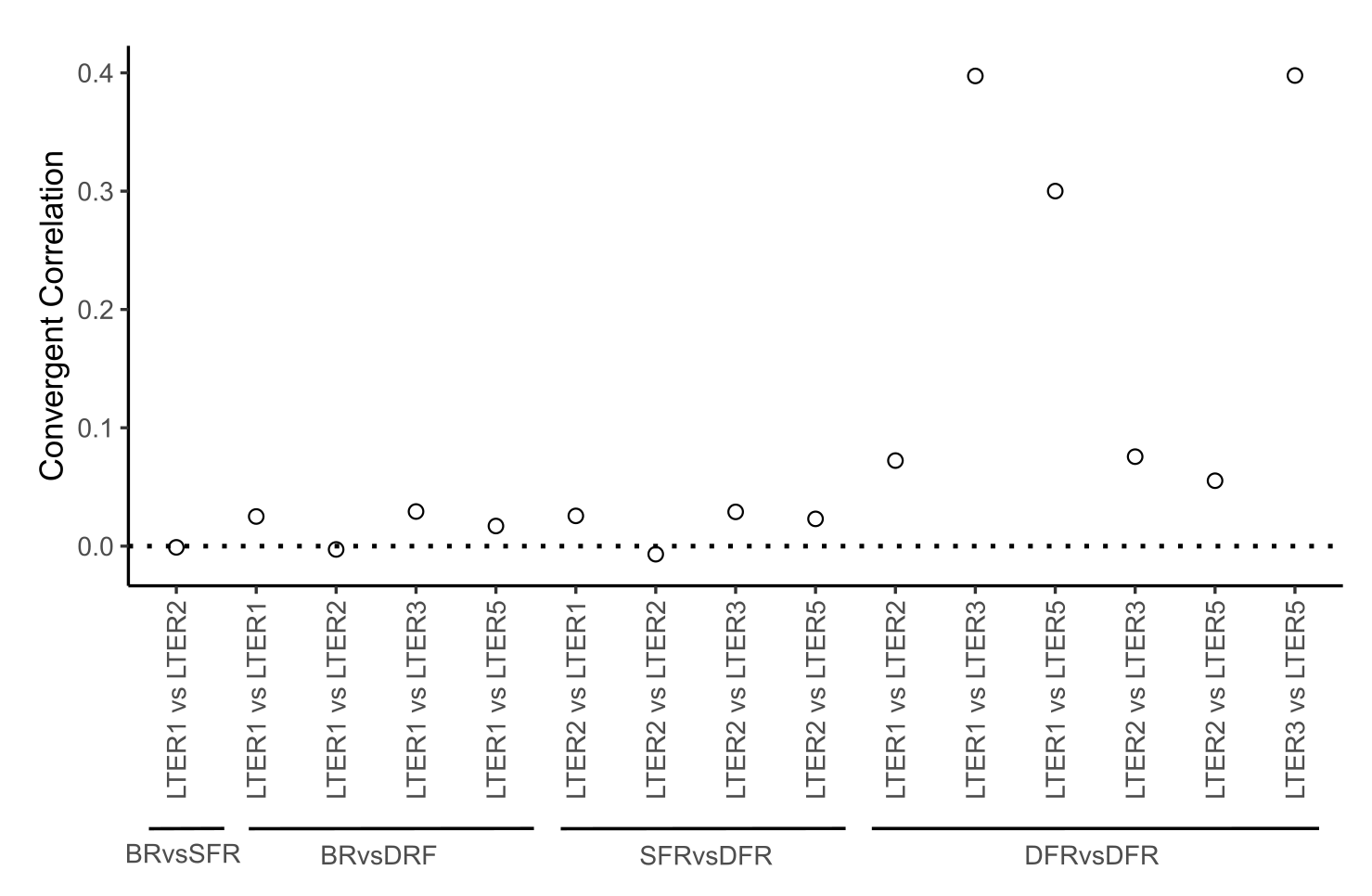

Supplemental Figure 20. Convergent correlation statistic for allele frequency shifts between pre- and post- mortality timepoints using only the deep forereef bleaching survival PGS (Fig. 3C). BR: Backreef; SFR: Shallow forereef; DFR: Deep forereef.

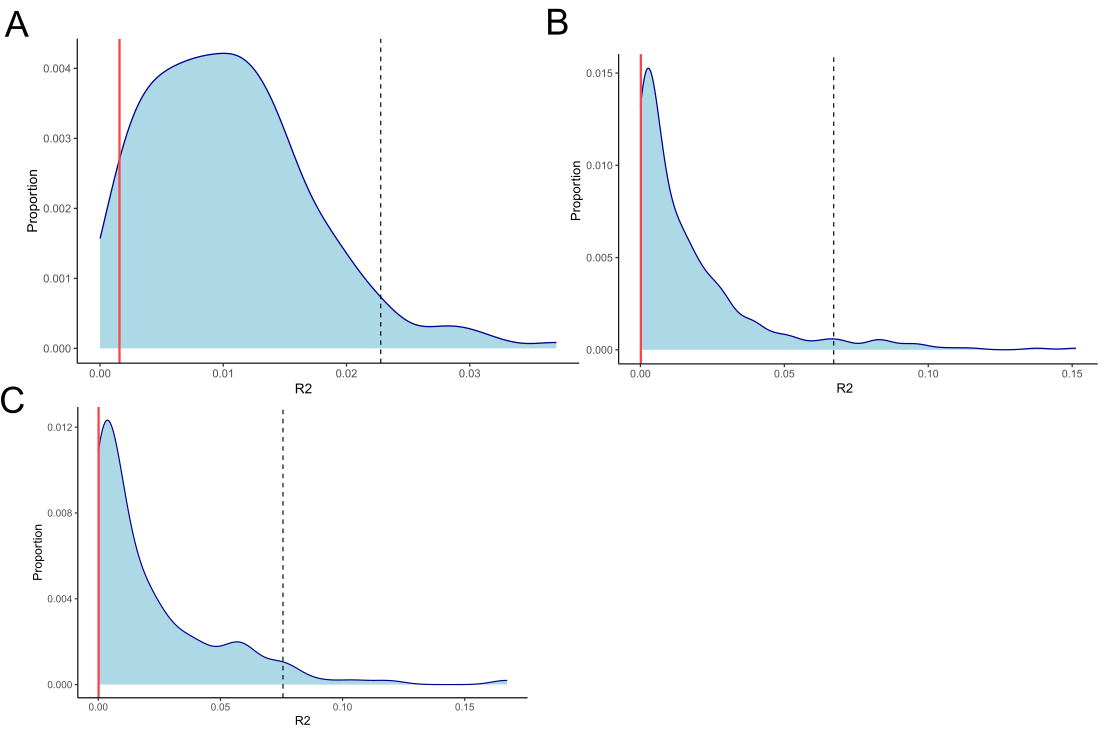

Supplemental Figure 21. Predictive power (R^2^) for backreef bleaching survival PGS for the 500 sets of randomly selected loci (blue distribution) and for the loci passing *p*-value A) < 0.01 B) < 0.001 and C) < 0.0001 thresholds for the deep forereef bleaching survival GWAS. Dotted line shows 95^th^ percentile for the null distribution and red line shows R^2^ for the real PGS.

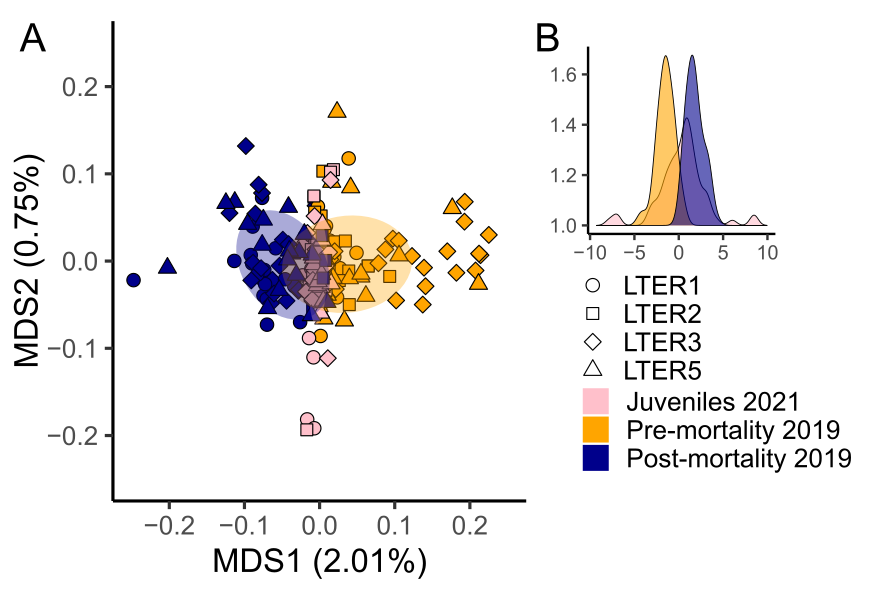

Supplemental Figure 22. MDS (A) and DAPC (B) plots based on genetic covariance matrices from the bleaching survival GWAS outlier loci only, demonstrating the intermediate state of juvenile genotypes.

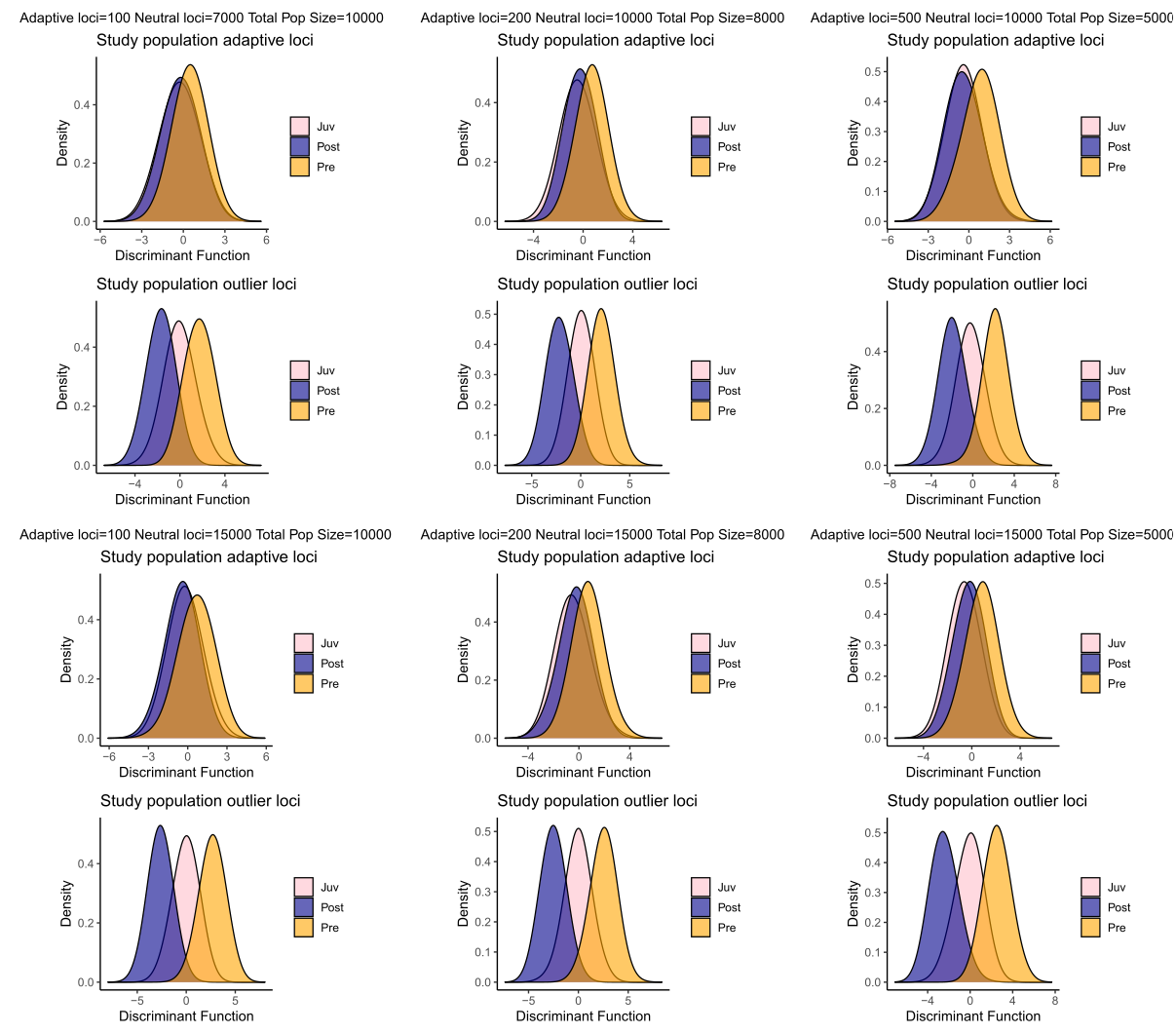

Supplemental Figure 23. Intermediate distribution from juveniles as a function of sampling bias. DAPCs for simulated data across different iterations of adaptive loci (*i.e.*, adaptive loci in the entire population), neutral loci (*i.e.*, neutral loci in the entire population) and population size (*i.e.*, population size of the entire population) examining both adaptive loci (top; true adaptive loci within outlier loci identified by GWAS) and outlier loci (bottom; all loci identified by GWAS).

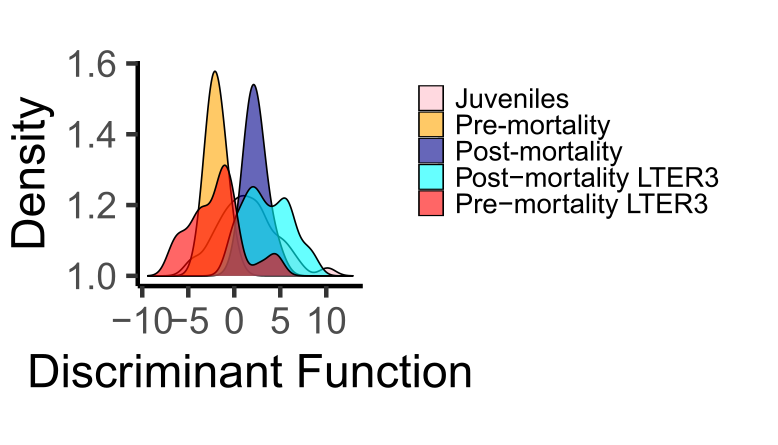
Supplemental Figure 24. DAPC plots based on genetic covariance matrices from the deep forereef bleaching survival GWAS outlier loci plotting LTER3 pre- and post-mortality separately.

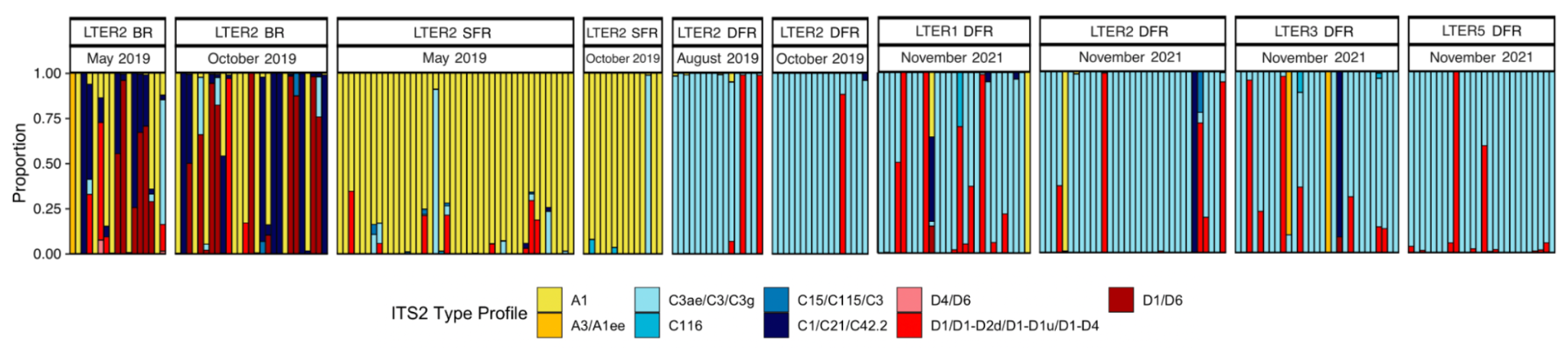

Supplemental Figure 25. Proportion of collapsed ITS2 type profiles from ITS2 metabarcoding data analyzed using SymPortal. Samples include those collected in 2019 at the backreef, shallow forereef, and deep forereef at LTER 2, published in Leinbach et al. (2023), and juveniles collected in 2021 from deep forereef sites (LTER 1, LTER 2, LTER 3, LTER 5). Colored bars correspond to relative abundances of dominant profile types in an individual colony.

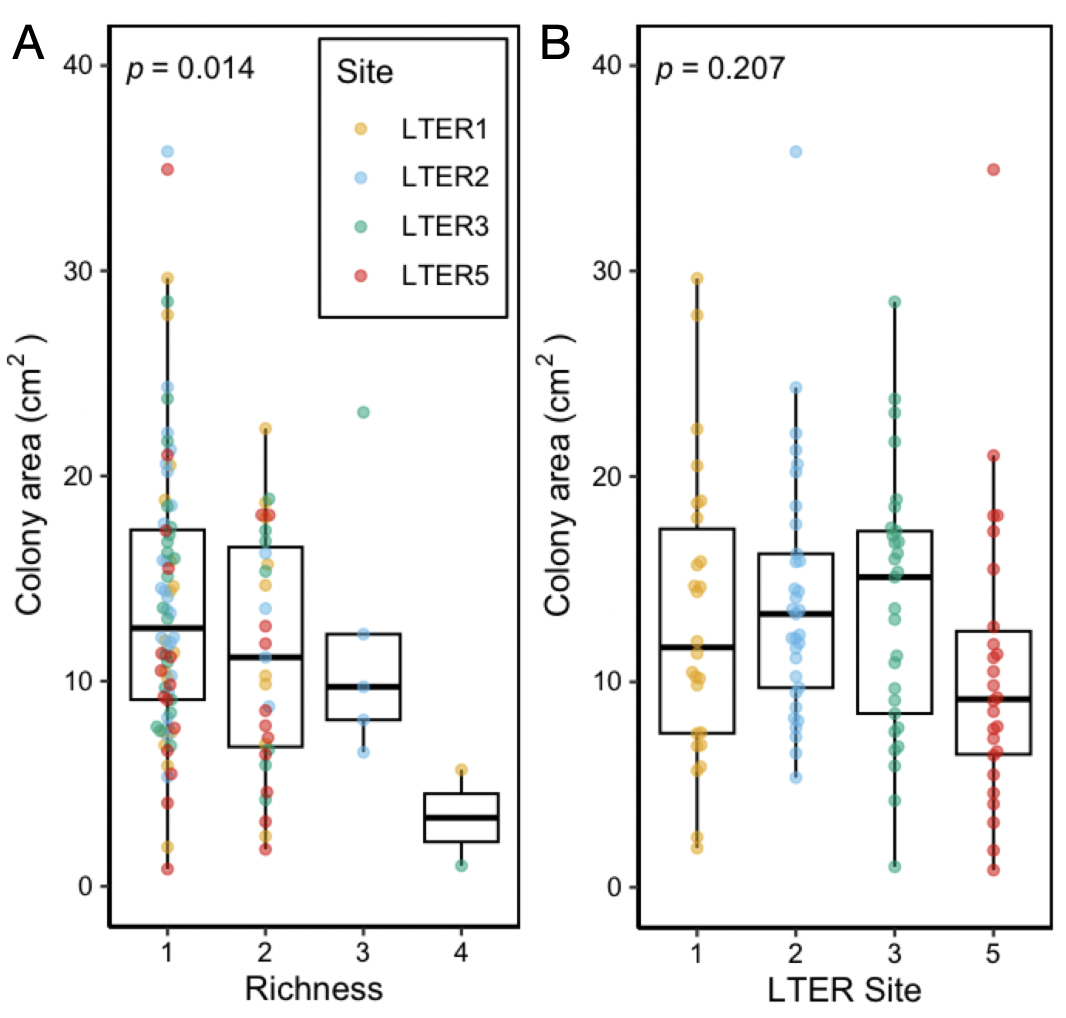

Supplemental Figure 26. Comparisons of juvenile colony area by richness and site (2021 juvenile samples at deep forereef sites only, N = 114). A) Colony area across ITS2 type profile richness. B) Colony area compared between sites. Each dot represents an individual colony.

Supplemental Figure 27. Non-metric multidimensional scaling (NMDS) plot of between sample symbiont community structure (LTER 2 deep forereef only) in 2019 and 2021 based on collapsed ITS2 type profiles.
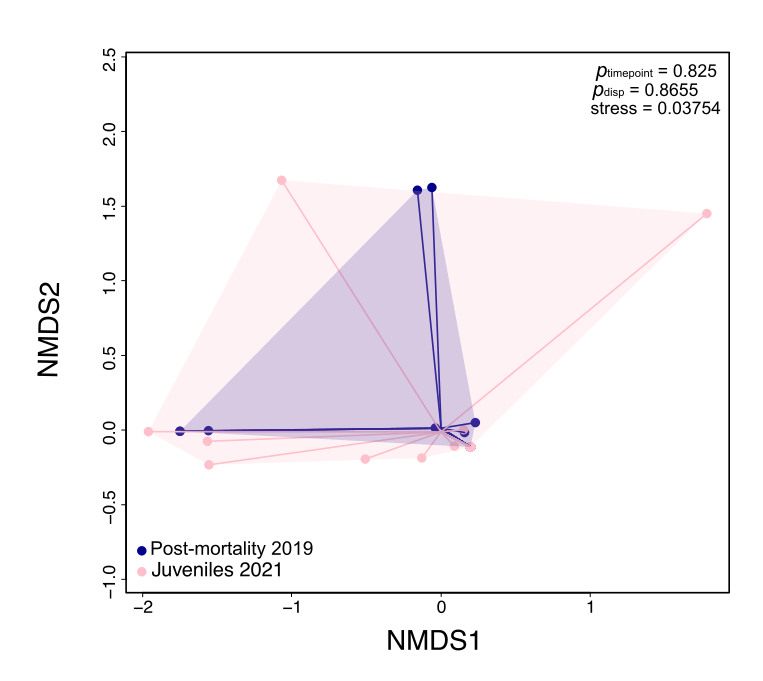

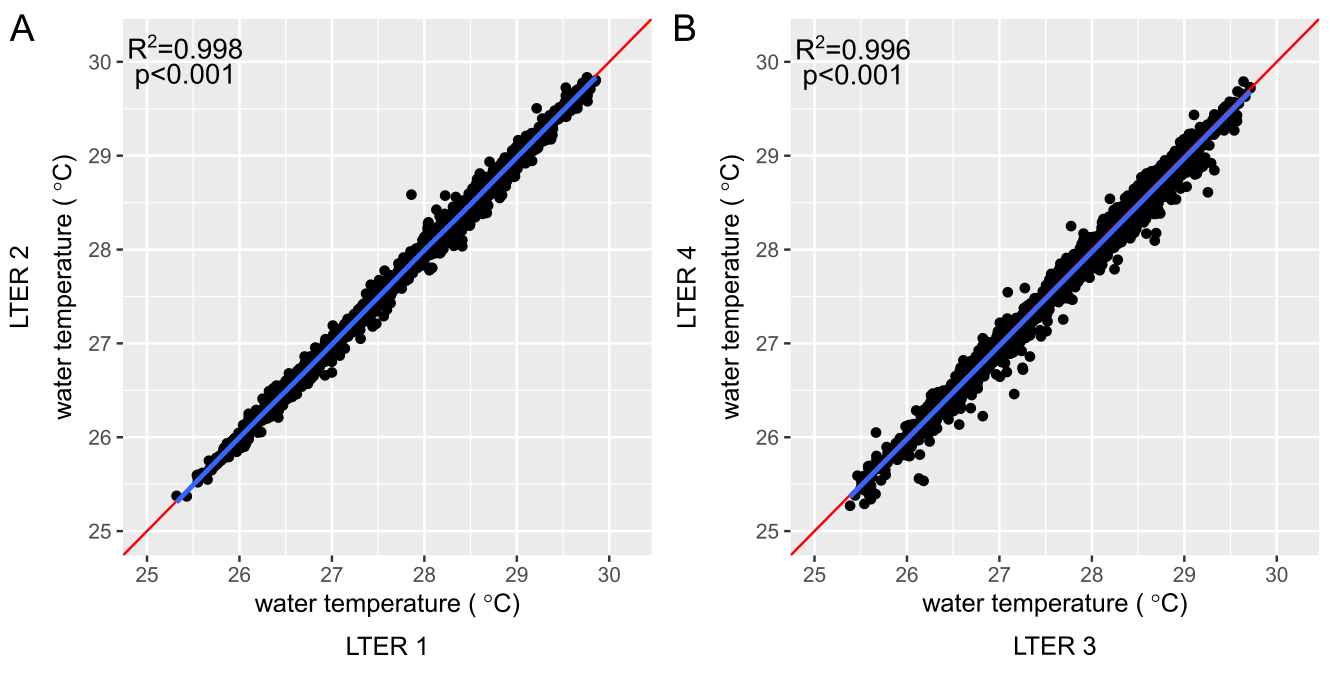

Supplemental Figure 28. Correlation between median daily water temperatures for substituted thermistors for Fig. 1 during periods of available overlapping data from May 2005 to August 2021 for A) North side and B) East side of Mo’orea.

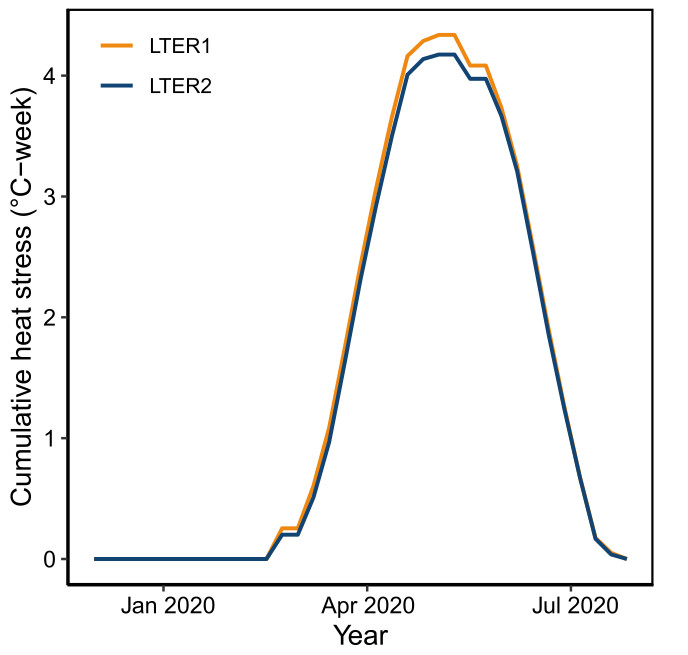

Supplemental Figure 29. Comparison of cumulative heat stress calculated for available overlapping data (*i.e.*, 2020) for substituted thermistors in Fig. 1 during periods of available overlapping data.

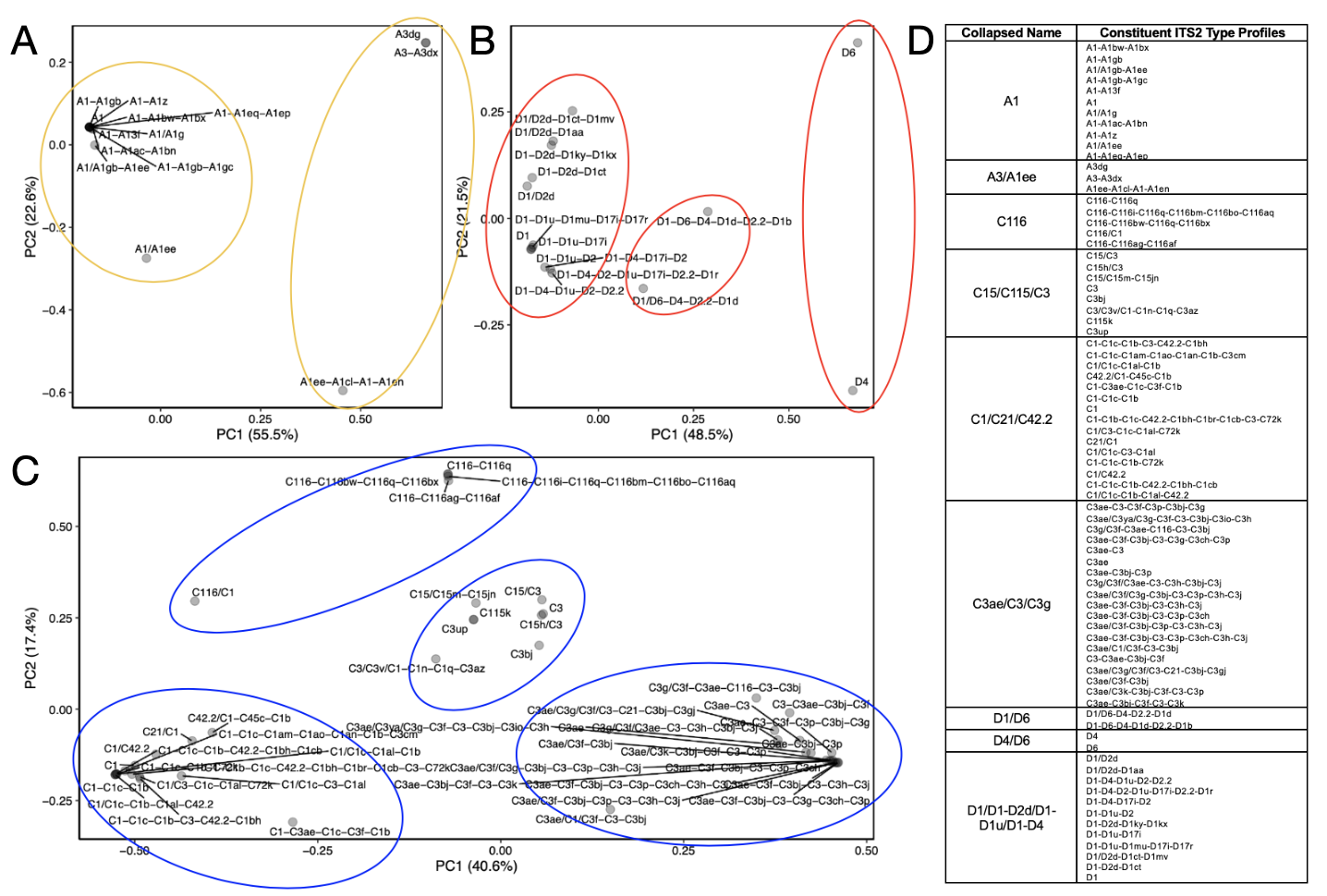

Supplemental Figure 30. Collapsed Symbiodiniaceae ITS2 type profiles of juveniles collected at deep forereef sites in 2021. PCA was conducted based on Bray-Curtis distances provided by SymPortal between A) *Symbiodinium*, B) *Durusdinium*, and C) *Cladocopium* profiles. Elliptical shapes were drawn around profiles based on similarity and/or their distribution along PC1. D) Table of nine collapsed ITS2 type profiles and their constituent ITS2 type profiles (78 profiles total). Collapsed profile names are derived from the dominant defining intragenomic variant (DIV) found within each constituent ITS2 type profile, with slashes separating DIVs by genotype.

x

Supplemental Table 1. Sample and site information for bleaching surveys, WGS and ITS2 sequencing.

| **LTER** | **Reef Habitat** | **Collection Timepoint** | **Lat** | **Long** | **Temp data source** | **Bleaching info (N)** | **WGS data (N) (clones removed)** | **ITS2 (N)** | **Depth (m)** |
| --- | --- | --- | --- | --- | --- | --- | --- | --- | --- |
| LTER 1 | Deep Forereef | May 2019 | -17.475929 | -149.8398389 | LTER 0 | 50 | 21 | 0 | 10 |
| LTER 1 | Deep Forereef | October 2019 | -17.475929 | -149.8398389 | LTER 0 | *NA* | 21 | 0 | 10 |
| LTER 1 | Deep Forereef | November 2021 | -17.475929 | -149.8398389 | LTER 0 | *NA* | 21 | 27 | 10 |
| LTER 1 | Shallow Forereef | May 2019 | -17.47628717 | -149.8397745 | LTER 0 | 52 | 16 | 0 | 3-5 |
| LTER 1 | Backreef | May 2019 | -17.4798382 | -149.8400535 | None | 25 | 19 | 0 | 1-3 |
| LTER 1 | Backreef | October 2019 | -17.4798382 | -149.8400535 | None | *NA* | 19 | 0 | 1-3 |
| LTER 2 | Deep Forereef | August 2019 | -17.473111 | -149.817639 | LTER 0 | 66 | 0 | 16 | 10 |
| LTER 2 | Deep Forereef | May 2019 | -17.473111 | -149.817639 | LTER 0 | 65 | 18 | 0 | 10 |
| LTER 2 | Deep Forereef | October 2019 | -17.473111 | -149.817639 | LTER 0 | *NA* | 14 | 17 | 14 |
| LTER 2 | Deep Forereef | November 2021 | -17.473111 | -149.817639 | LTER 0 | *NA* | 19 | 33 | 10 |
| LTER 2 | Shallow Forereef | May 2019 | -17.4738068795 | -149.8172635 | LTER 0 | 63 | 23 | 42 | 3-5 |
| LTER 2 | Shallow Forereef | October 2019 | -17.4738068795 | -149.8172635 | LTER 0 | *NA* | 13 | 14 | 3-5 |
| LTER 2 | Backreef | May 2019 | -17.476204 | -149.813934 | None | 25 | 0 | 17 | 1-3 |
| LTER2 | Backreef | October 2019 | -17.476204 | -149.813934 | None | *NA* | 0 | 27 | 1-3 |
| LTER 3 | Deep Forereef | May 2019 | -17.51547255 | -149.7620307 | LTER 4 | 72 | 28 | 0 | 10 |
| LTER 3 | Deep Forereef | October 2019 | -17.51547255 | -149.7620307 | LTER 4 | *NA* | 19 | 0 | 10 |
| LTER 3 | Deep Forereef | November 2021 | -17.51547255 | -149.7620307 | LTER 4 | *NA* | 21 | 29 | 10 |
| LTER 5 | Deep Forereef | May 2019 | -17.57837964 | -149.8758341 | LTER 5 | 100 | 14 | 0 | 10 |
| LTER 5 | Deep Forereef | October 2019 | -17.57837964 | -149.8758341 | LTER 5 | *NA* | 19 | 0 | 10 |
| LTER 5 | Deep Forereef | November 2021 | -17.57837964 | -149.8758341 | LTER 5 | *NA* | 16 | 26 | 10 |
| LTER 5 | Backreef | May 2019 | 17.57552605 | -149.876156 | LTER 5 | 19 | 0 | 0 | 1-3 |

Supplemental Table 2. RDA model using a combination of explanatory and conditional variables with the response variable as host genetic variation.

| **Explanatory variable** | **Conditional variables** | **R^2^** | **p-value** |
| --- | --- | --- | --- |
| Dominated by *Symbiodinium* (Yes/No) | Depth + LTER1 + LTER2 + LTER3 + LTER5 | <0.01 | 0.614 |
| Dominated by *Cladocopium* (Yes/No) | Depth + LTER1 + LTER2 + LTER3 + LTER5 | <0.01 | 0.648 |
| Dominated by *Durusdinium* (Yes/No) | Depth + LTER1 + LTER2 + LTER3 + LTER5 | <0.01 | 0.434 |

References

Edmunds, P., & Moorea Coral Reef LTER. (2022). MCR LTER: Coral Reef: Long-term Population and Community Dynamics: Corals, ongoing since 2005. In *LTER Network Member Node.* https://pasta.lternet.edu/package/metadata/eml/knb-lter-mcr/4/39

Fifer, J. E., Yasuda, N., Yamakita, T., Bove, C. B., & Davies, S. W. (2022). Genetic divergence and range expansion in a western North Pacific coral. *Science of The Total Environment*, *813*, 152423. https://doi.org/10.1016/j.scitotenv.2021.152423

Leinbach, S. E., Speare, K. E., & Strader, M. E. (2023). Reef habitats structure symbiotic microalgal assemblages in corals and contribute to differential heat stress responses. *Coral Reefs*, *42*(1), 205–217. https://doi.org/10.1007/s00338-022-02316-w

Rose, N. H., Bay, R. A., Morikawa, M. K., Thomas, L., Sheets, E. A., & Palumbi, S. R. (2021). Genomic analysis of distinct bleaching tolerances among cryptic coral species. *Proceedings of the Royal Society B: Biological Sciences*, *288*(1960). https://doi.org/10.1098/rspb.2021.0678
